# MicrobeRT: A Framework and Suite of Fine-tuned Language Models for Genomic Sequences

**DOI:** 10.64898/2026.11.03.763945

**Authors:** Vijay Narayan, Brian Merritt, Sanjeevani Choudhery, Daniel Jimenez, Stanley Ta, Daniel S. Berman, Jeremy Ratcliff, Molly E. Gallagher

## Abstract

Accurate and reproducible metagenomic analysis is essential for effective infectious disease surveillance. Targeted diagnostics such as culturing, antigen assays, and multiplex PCR panels identify only predefined organisms and can fail to identify atypical or novel infections. Shotgun metagenomics offers hypothesis-free detection, but adoption is limited by workflow complexity, dependence on mutable reference databases, and high computational cost.

To combat these challenges, we introduce MicrobeRT, an integrated suite of genomic language models fine-tuned to classify metagenomic sequencing reads across three tasks: (i) species level taxonomic identification, (ii) antibiotic resistance gene detection, and (iii) pathogenicity assessment. The framework builds on three pre-trained genomic language models—Nucleotide Transformer [1], DNABERT [2, 3], and HyenaDNA [4]—leveraging their large-scale genomic pre-training, adaptability to downstream tasks, and, in the case of HyenaDNA, ability to capture long-range sequence dependencies. The models are fine-tuned on large, curated datasets spanning microbial, viral, fungal, and human genomes, including more than 500,000 genomes and 250,000 antibiotic resistance gene sequences.

To evaluate the performance of our fine-tuned genomic language models across classification tasks, we developed a benchmarking framework that compares MicrobeRT against state-of-the-art (SOTA) bioinformatics tools. Across both simulated and empirical datasets, the models achieve strong performance while reducing computational overhead. MicrobeRT is released as open-source software, enabling further fine-tuning and reproducible integration of alternative models and classifiers. We also demonstrate seamless, drop-in integration by incorporating our fine-tuned taxonomic classification models as an optional add-on within the open-source TaxTriage pipeline [5]. MicrobeRT demonstrates the feasibility of three clinically relevant multi-label analyses, offering a practical path to scalable, reproducible metagenomic diagnostics and enabling broader deployment of AI/ML-enhanced infectious disease testing.

---

Accurate and reproducible pathogen detection is vital for effective infectious disease surveillance. Traditional diagnostic methods, such as culture-based assays, antigen detection, and polymerase chain reaction (PCR) panels that have long been central to clinical microbiology fail when pathogens are atypical or novel, making serial testing costly and ineffective. Targeted methods such as these are also constrained to a finite number of organisms. Multiplex assays expand diagnostic coverage, but as of 2026, even the largest FDA-cleared PCR panels detect only 43 pathogens at one time [6].

Untargeted sequencing approaches such as shotgun metagenomics offer a powerful alternative. Shotgun metagenomics allows for hypothesis-free identification of both known and highly divergent or novel pathogens. Beyond pathogen detection, metagenomics also enables downstream analyses such as taxonomic classification and prediction of antimicrobial resistance. Numerous cloud-based platforms such as BugSeq [7] and CZID [8] have been developed to make these analyses accessible. However, widespread adoption of metagenomic diagnostics has been limited by reproducibility challenges and workflow complexity. Each step of the analytic workflow, from read cleaning and taxonomic classification to alignment and assembly, relies on different software and reference databases that can have substitutions, leading to variability in results, reducing reproducibility. Expanding these databases can improve sensitivity and specificity, but it comes at higher time and computational costs. Additionally, reliance on external infrastructure introduces maintenance overhead and dependency risks.

To address these challenges, new analytic approaches are needed that reduce the dependence on reference databases while improving speed, scalability, and reproducibility. Recent advances in language models offer a promising path forward. Like words in natural languages, nucleotide (and protein) sequences encode biological information through ordered sequences of discrete symbols. Genomic language models (gLMs) trained on large databases of genomic sequences have demonstrated high proficiency in predicting functional elements such as promoters, enhancers, and splice sites (see [9] for a review). It has also been demonstrated that fine-tuning can adapt large pre-trained models for specific classification applications [1]. However, gLMs have not been developed specifically for species-level classification of microbial sequences, nor do current approaches unify multiple diagnostic metagenomic multi-label classification tasks into a single reproducible framework.

Here we introduce MicrobeRT, an integrated framework and suite of genomic language models fine-tuned for three distinct tasks: taxonomic classification, antimicrobial resistance gene detection, and pathogen detection. MicrobeRT builds on the complementary strengths of three open-source foundation model architectures: Nucleotide Transformer [1], DNABERT-2 [2] and DNABERT-S [3], and HyenaDNA [4]. MicrobeRT models are fine-tuned on large, curated datasets spanning over 500,000 microbial, viral, and fungal genomes and over 250,000 annotated antimicrobial resistance (AMR) sequences. The fine-tuning and evaluation framework developed in this work is readily configurable for diverse downstream classification tasks. The codebase is publicly available on GitHub (https://github.com/jhuapl-bio/microbert), and all fine-tuned models are available on Hugging Face (https://huggingface.co/jhuapl-bio/microbert).

We benchmarked the performance of our fine-tuned gLMs against SOTA database-driven and machine learning methods including Kraken2 [10], SeqScreen [11], CZID [8], BERTax [12], DeepARG [13], DeePaC [14], and Lyra [15]. This was done using both simulated reads and real-world Illumina and Oxford Nanopore sequencing data for three core bioinformatics tasks: taxonomic classification, antimicrobial resistance (AMR) gene detection, and pathogenicity prediction. For taxonomic classification, MicrobeRT models performed best for taxa and genes well represented in the fine-tuning data, with notable improvements for clinically relevant organisms such as *Bordetella pertussis*. In contrast, database-driven methods such as Kraken2 retained advantages for sparsely represented taxa. For AMR prediction, the fine-tuned NT and HyenaDNA models outperformed DeepARG across binary and multi-label settings and remained robust to truncated and flanking sequence contexts, highlighting their ability to recognize resistance signals beyond exact sequence matches. For pathogenicity classification, performance was task-dependent: Lyra excelled on fungal datasets, Hye-naDNA on bacterial data, and DeePaC [14] performed best on viral sequences. Finally, we integrated the taxonomic classification capability of MicrobeRT into the TaxTriage metagenomic workflow [5], enabling practical deployment alongside existing tools and demonstrating that sequence-based genomic language models can complement traditional pipelines by providing enhanced sensitivity to novel or divergent organisms where reference-based methods often fail.

By unifying taxonomic classification, AMR detection, and pathogen discrimination in a single reproducible framework not dependent on an external reference database, MicrobeRT enhances reproducibility across sequencing platforms and reduces computational overhead. MicrobeRT is integrated into the open-source TaxTriage pipeline as an optional complement to traditional classifiers such as Kraken2, enabling direct comparison within established bioinformatics workflows. More broadly, this work illustrates that foundation models for genomic sequences can be adapted into practical, scalable tools for infectious disease surveillance and public health response, particularly in environments involving novel or divergent organisms where reference-based methods may be limited.

## Results

### Leveraging complementary genomic foundation models for metagenomic classification

Recent advances in genomic language modeling have led to a rapid increase in publicly available pretrained models. In this work, we leverage three such model architectures—Nucleotide Transformer (NT) [1], DNABERT [2, 3], and HyenaDNA [4]—and fine-tune them for three tasks: taxonomic classification, AMR gene detection, and pathogen detection. We refer to the resulting framework and suite of fine-tuned models collectively as “MicrobeRT”. These models span both transformer and non-transformer architectures and differ in tokenization strategies, scale, and sequence-length capacity. For each architecture, we evaluate multiple variants: NT (50M, 100M, and 250M parameters), DNABERT-2 (117M parameters), and HyenaDNA models supporting sequence lengths of 32k, 150k, 320k, 450k, and 1M tokens. For each task, we appended a task-specific linear classification head and fine-tuned the model end-to-end. Each model accepts a nucleotide sequence as input and outputs a binary or multi-label classification prediction. Because these architectures differ in their ability to represent short- and long-range sequence context, we anticipated performance to vary across tasks and read-length regimes. We benchmarked MicrobeRT against database-driven and task-specific baselines across simulated and empirical datasets.

### Complementation of reference-based taxonomic classifiers, with strongest gains on long reads and well-supported taxa

We benchmarked MicrobeRT for taxonomic classification against the SOTA reference-based classifiers Kraken2 [10], Centrifuge [16], Kaiju [17], DIAMOND [18], and SeqScreen [11]. We also benchmarked against BERTax [19], a transformer-based model trained from scratch for hierarchical classification across superkingdom, phylum, and genus. Taxonomic data for training were curated as described in our Materials and Methods section, and a 1% hold-out set was reserved for testing. Across benchmarks, performance depended strongly on evaluated sequence length and taxon representation in the fine-tuning data. HyenaDNA achieved the strongest performance on long fragments (500–1,750 bp), consistent with its greater context capacity, while NT and DNABERT-2 were more competitive on short fragments (50-300 bp) (see Figures 2, 3).

Across read-length regimes, MicrobeRT models performed best for taxa that were well represented during fine-tuning, including clinically relevant organisms such as *Bordetella pertussis*, whereas Kraken2 retained an advantage for sparsely represented taxa, as seen in Figure 1. To evaluate taxonomic classification, sequences were derived from representative bacterial, fungal, viral, and archaeal RefSeq genomes, subset to the most common and relevant pathogens consisting of almost 1000 unique species. In this setting, Kraken2 outperformed the fine-tuned NT 50M model frequently at short-read species classification and fine-tuned HyenaDNA 1M model at long-read species classification (Figure 1A). However, MicrobeRT performance improved with greater training support per taxon, and for highly represented species, the fine-tuned models exceeded the performance of the reference-based classifiers. Across the broader RefSeq benchmark, it is seen that taxa with the strongest representation in the fine-tuning set showed markedly higher F1 scores than SOTA classifiers (Figure 2). Notably, the 10 species with the greatest training support all showed substantial gains relative to the baseline methods (Figure 2B). These results suggest that, for short reads, MicrobeRT is most effective when sufficient taxon-specific training signal is available, and that it may benefit further from targeted fine-tuning on organisms of clinical interest.

**Figure 1:**
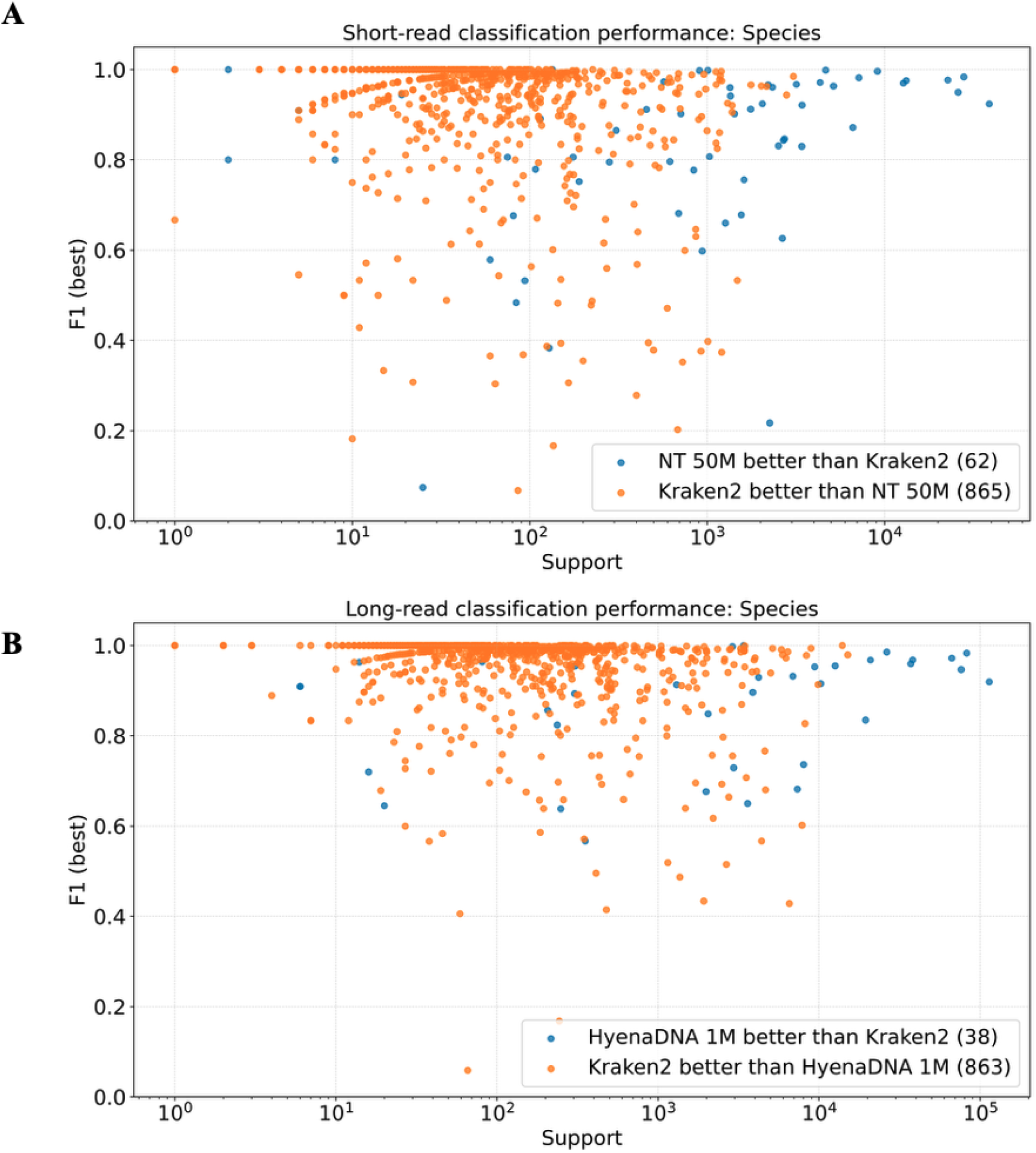
Performance in species classification (F1 score) vs. support (number of reads included in the test data, which is directly correlated with support in the training data) of the fine-tuned MicrobeRT models and Kraken2 for (A) short-read (<300 bp) sequence classification with Nucleotide Transformer 50M and (B) long-read sequence classification with HyenaDNA 1M.

**Figure 2:**
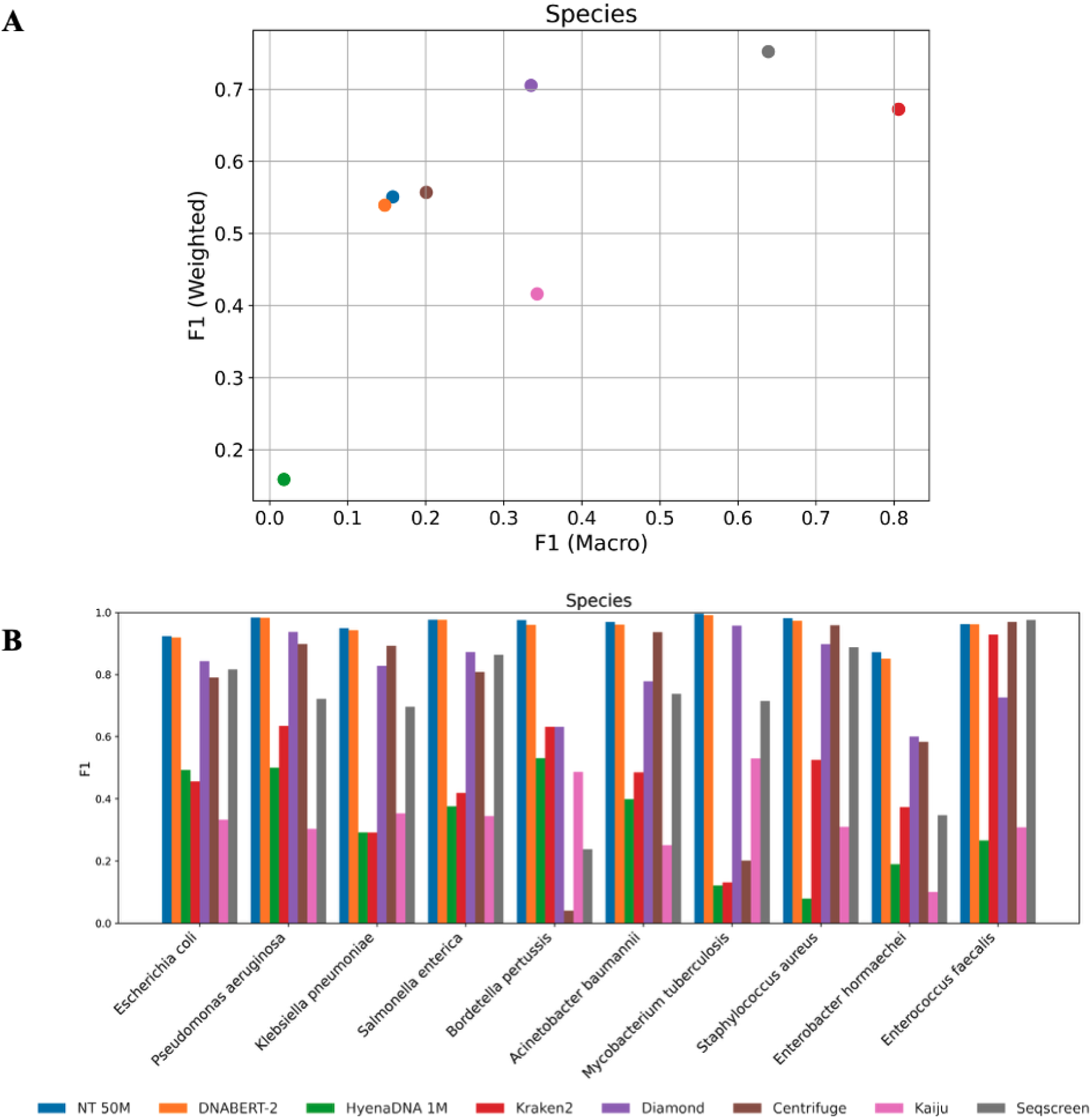
F1 scores for short-read (*<*300 bp) sequences, across species (A) and for each of the 10 species with the most support in the fine-tuning data (B).

For phylum and genus-level classification on sequences with taxonomic labels derived from the BERTax benchmark dataset, our fine-tuned HyenaDNA model exceeded BERTax—the closest sequence-based baseline—across macro-, micro-, and weighted F1 scores, as well as mean reciprocal rank (MRR) (Figure 4). Notably, BERTax does not support species-level classification, and thus no direct comparison was performed at that resolution. As in the short-read setting, long-read performance was highest for taxa that were well represented in the fine-tuning data and lower for sparsely represented taxa (Figure 3).

**Figure 3:**
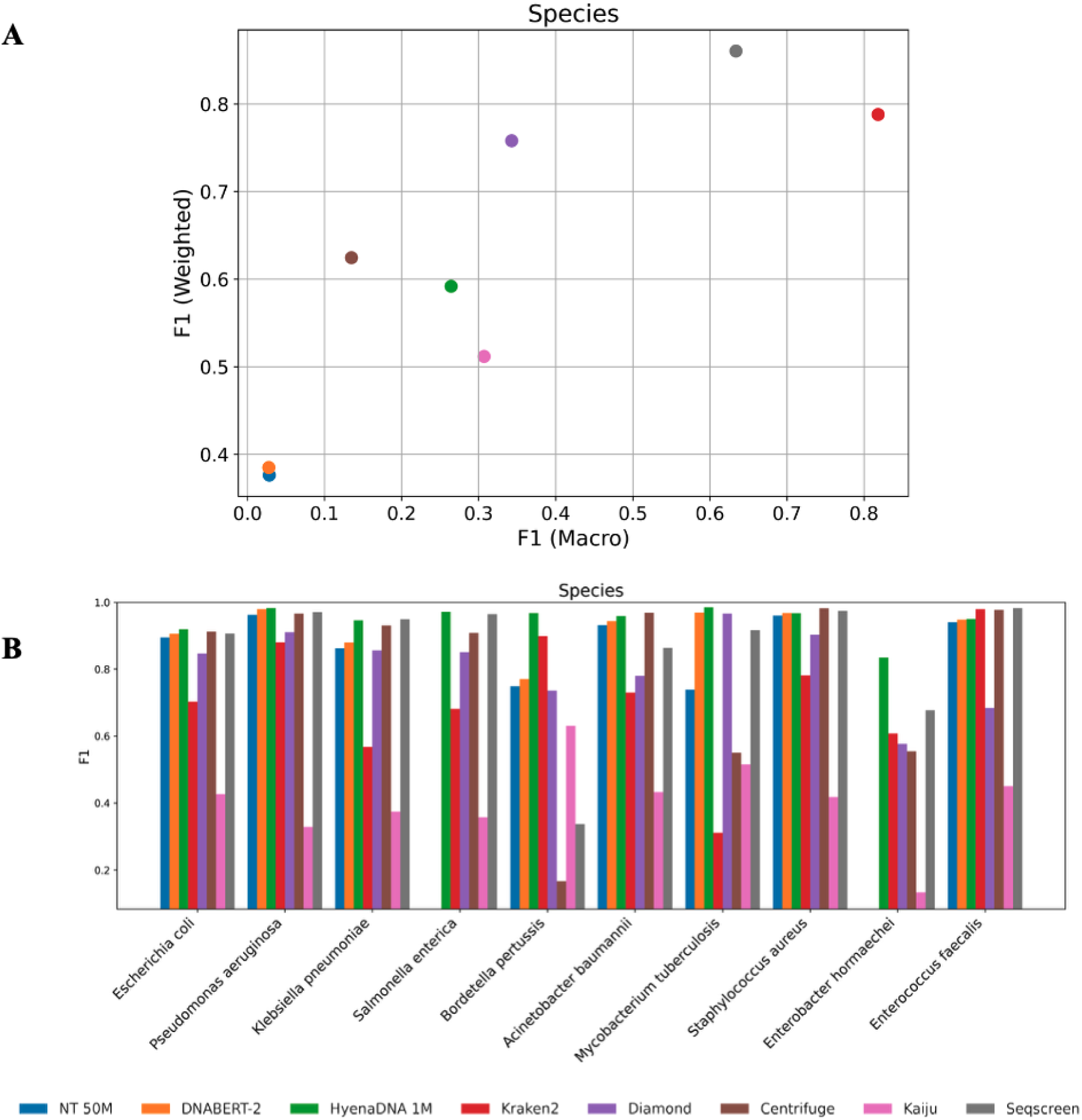
F1 scores for long-read (500–1750 bp) sequences, across species (A) and for each of the 10 species with the most support in the fine-tuning data (B).

**Figure 4:**
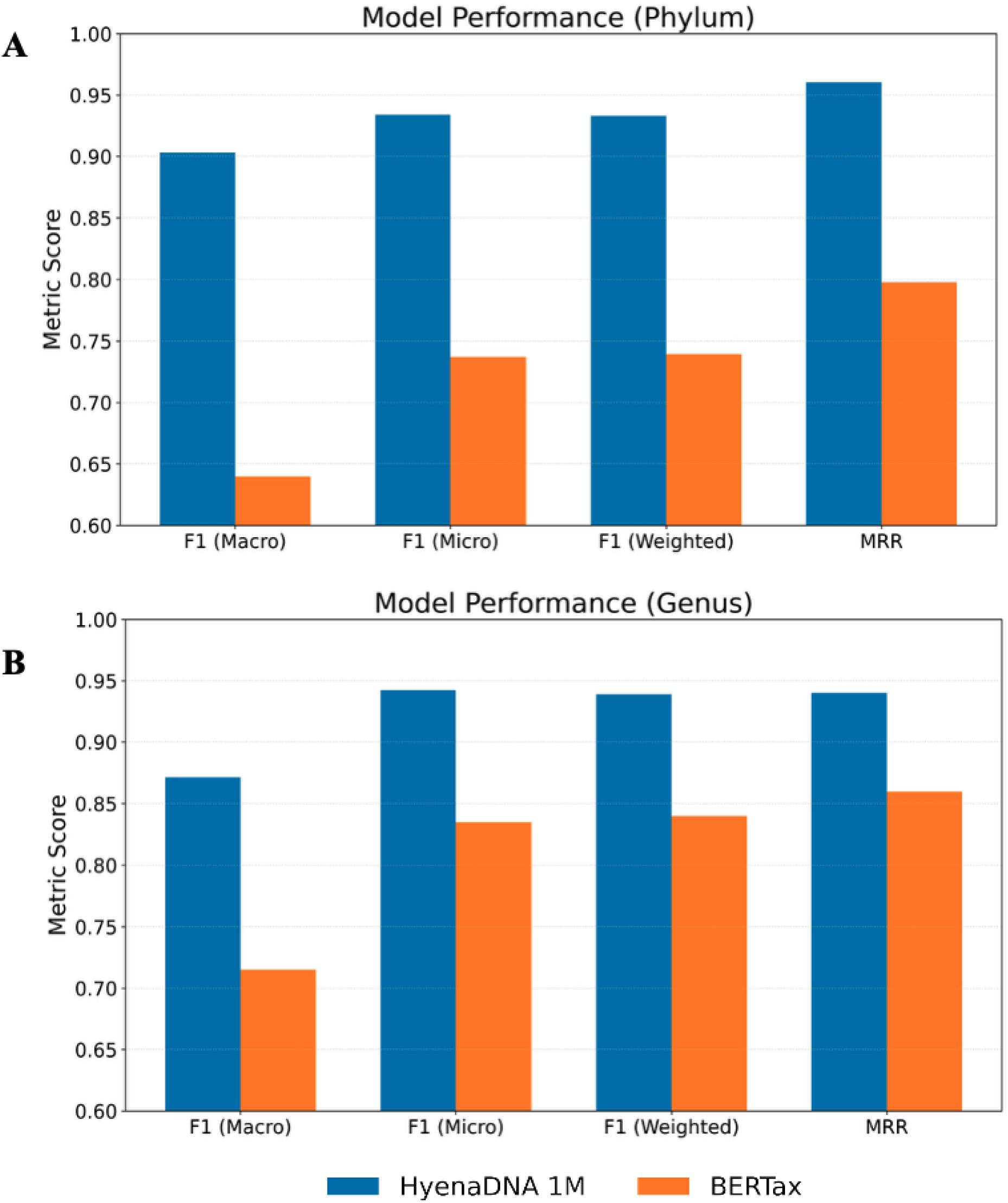
Phylum (A) and Genus (B) level classification performance of the fine-tuned HyenaDNA 1M model compared with BERTax. HyenaDNA outperformed BERTax across all evaluation metrics on the published BERTax test set [19].

Overall, HyenaDNA provided the best taxonomic classification performance on long fragments, whereas NT and DNABERT-2 performed best on shorter reads. MicrobeRT models notably out-performed all comparator methods for selected species of interest, including *Bordetella pertussis*, but Kraken2 remained superior in most low-support settings. Taken together, these findings show that MicrobeRT complements rather than replaces k-mer and alignment-based classifiers, with its clearest advantages emerging for long reads and for taxa that are well supported during fine-tuning.

### Detection of AMR signals across truncation and flanking-context perturbations

Accurate detection of AMR gene presence is clinically important because resistance calls can help inform treatment decisions, support antimicrobial selection and management, and prioritize downstream confirmatory testing. We benchmarked MicrobeRT against SOTA AMR classification approaches, including DeepARG [13], a deep learning based sequence classifier for antimicrobial resistance genes, and CZID [8], a cloud-based metagenomic analysis platform. To determine whether MicrobeRT identified AMR solely through exact sequence identity or could also capture resistance-associated signals beyond exact matches, we evaluated performance in both binary and multi-label classification settings on a 2% held-out test split of curated AMR gene sequences from the CARD and NCBI AMR Reference databases under five conditions: full-length AMR gene sequences, randomly truncated fragments (140–450 bp), and full-length AMR genes flanked on both ends by 50 bp, 100 bp, or randomly distributed lengths of 50–2,500 bp. These perturbations were designed to approximate the fragmented genes and heterogeneous sequence contexts encountered in metagenomic data.

For binary AMR classification tasks, our fine-tuned NT and HyenaDNA models showed the strongest performances and outperformed DeepARG across all evaluated test conditions (Figures 5). Lyra also exceeded DeepARG on the held-out test set but remained less competitive than the best-performing MicrobeRT models (Figure 5A). In contrast, the fine-tuned DNABERT-2 and DNABERT-S models performed poorly and often close to the majority class baseline metric, defined as the accuracy obtained by assigning every sequence to the most frequent class. Such a large performance gap between the DNABERT-based models and the NT and HyenaDNA models may be due to differences in model architecture, pre-training data, or tokenization, but conducting ablation studies to confirm this is beyond the scope of this work. Together, these findings indicated that the strongest MicrobeRT models can reliably distinguish AMR gene from non-AMR gene sequences even when AMR genes are truncated or embedded within additional flanking sequence.

**Figure 5:**
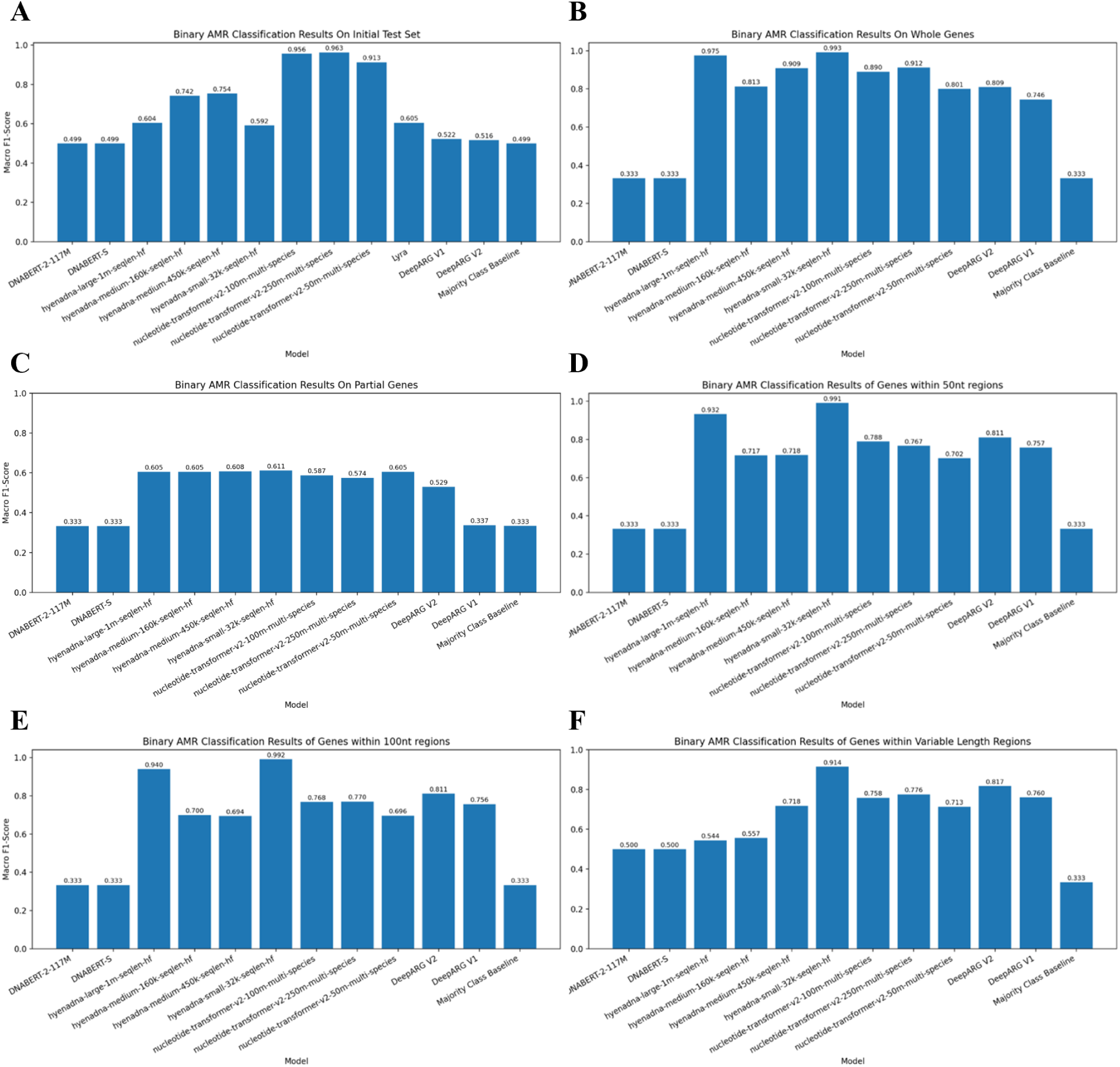
Macro F1 score of all models in predicting binary AMR classification for initial test set of the data (A), whole gene sequence data (B), partial gene sequence data (C), whole gene sequence data with an additional 50 nucleotides on either end (D), whole gene sequence data with an additional 100 nucleotides on either end (E), and whole gene sequence data with an additional random sequence length of 50-2500 nucleotides on either end (F).

For multi-label AMR classification, MicrobeRT again showed strong performance across all sequence conditions (Figure 6). Here, NT-based models, particularly NT-250M, performed best overall, with DNABERT-S consistently ranking next among the strongest models, both exceeding DeepARG across all test sets. Lyra again outperformed DeepARG on the held-out test set but did not match the best-performing genomic language models (Figure 6A). These findings suggest that the strongest MicrobeRT models can also preserve fine-grained resistance label assignment across diverse sequence contexts rather than simply detect AMR gene presence.

**Figure 6:**
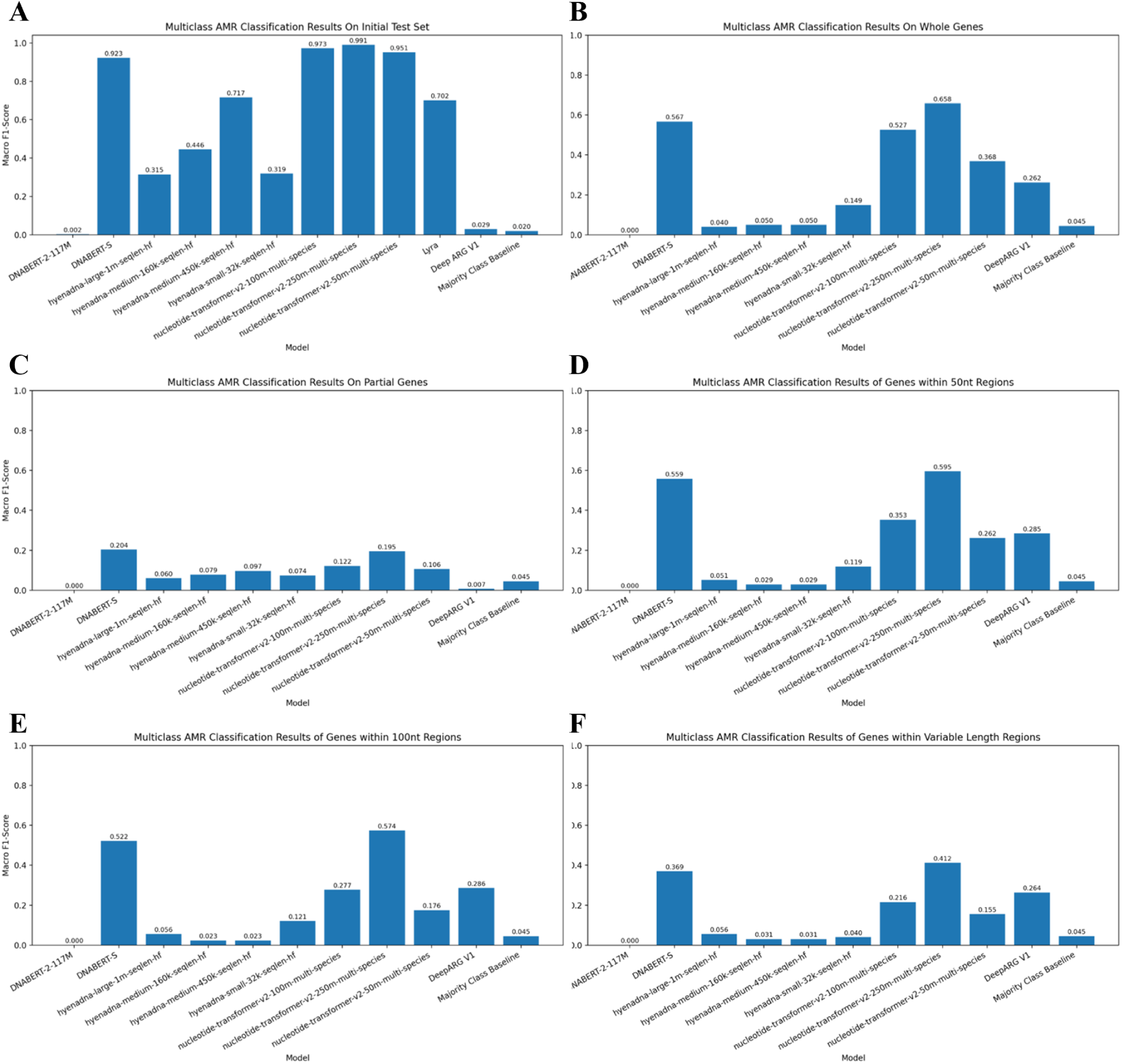
Macro F1 score of all models in predicting multi-label AMR classification for initial test set of the data (A), whole gene sequence data (B), partial gene sequence data (C), whole gene sequence data with an additional 50 nucleotides on either end (D), whole gene sequence data with an additional 100 nucleotides on either end (E), and whole gene sequence data with an additional random sequence length of 50-2500 nucleotides on either end (F).

As CZID’s AMR pipeline accepts only short-read Illumina data, it could not be directly bench-marked on these perturbed test sets. We therefore assessed CZID qualitatively using a representative run, which produces a summary table listing all detected AMR genes, their associated drug classes and mechanisms, and the number of mapped reads. Users can also access intermediate mapping files for detailed alignment statistics. Although this output is not directly comparable to the controlled evaluations used for MicrobeRT and DeepARG, it illustrates the type of AMR gene reporting provided by database-driven workflows and still provides performance information of CZID. Taken together, our evaluation demonstrates that MicrobeRT improves AMR detection across both binary and multi-label tasks and remains robust when those genes are fragmented or embedded within variable flanking sequence, supporting its use as a clinically relevant complement to existing AMR detection approaches.

### Pathogenicity classification reveals kingdom-specific strengths rather than a single dominant architecture

We next evaluated pathogenicity classification of MicrobeRT models against DeePaC [14], a deep-learning model based on reverse-complement neural networks that predicts pathogenic potential directly from DNA reads, and SeqScreen [11], a sequence-screening pipeline that combines alignment-based annotation with curated Functions of Sequences of Concern (FunSoCs) supported by ensemble machine learning. Because SeqScreen reports FunSoC assignments rather than direct binary pathogenicity labels, sequences assigned to any of the 32 FunSoCs were labeled as pathogenic for comparison.

For taxonomic and AMR classification, we primarily report performance using macro F1 score, which balances precision and recall while weighting each class equally. For binary pathogenicity classification, we instead use balanced accuracy, which averages the true positive and true negative rates. Unlike F1 score, which balances precision and recall, balanced accuracy gives equal weight to performance on pathogenic and non-pathogenic sequences, accounting for both false positive and false negative rates.

In contrast to taxonomy classification and AMR detection, pathogenicity classification showed stronger dataset-and kingdom-specific effects, indicating that this task was more heterogeneous and less uniformly captured by a single model family. DeePaC remained the strongest overall model, achieving the highest balanced accuracy on both viral datasets and consistently outperforming SeqScreen across the evaluated pathogen types (Figure 7). Among the MicrobeRT models, HyenaDNA-32k achieved the highest performance on the DeepSimulator bacterial dataset (See Materials and Methods-Pathogenicity data for details). Lyra performed best on DeePaC’s fungal dataset, with NT and HyenaDNA ranking second. NT-50M and HyenaDNA-32k performed near baseline on viral datasets, and DNABERT-2 remained near baseline across all pathogenicity tasks, consistent with its weaker performance in AMR classification.

**Figure 7:**
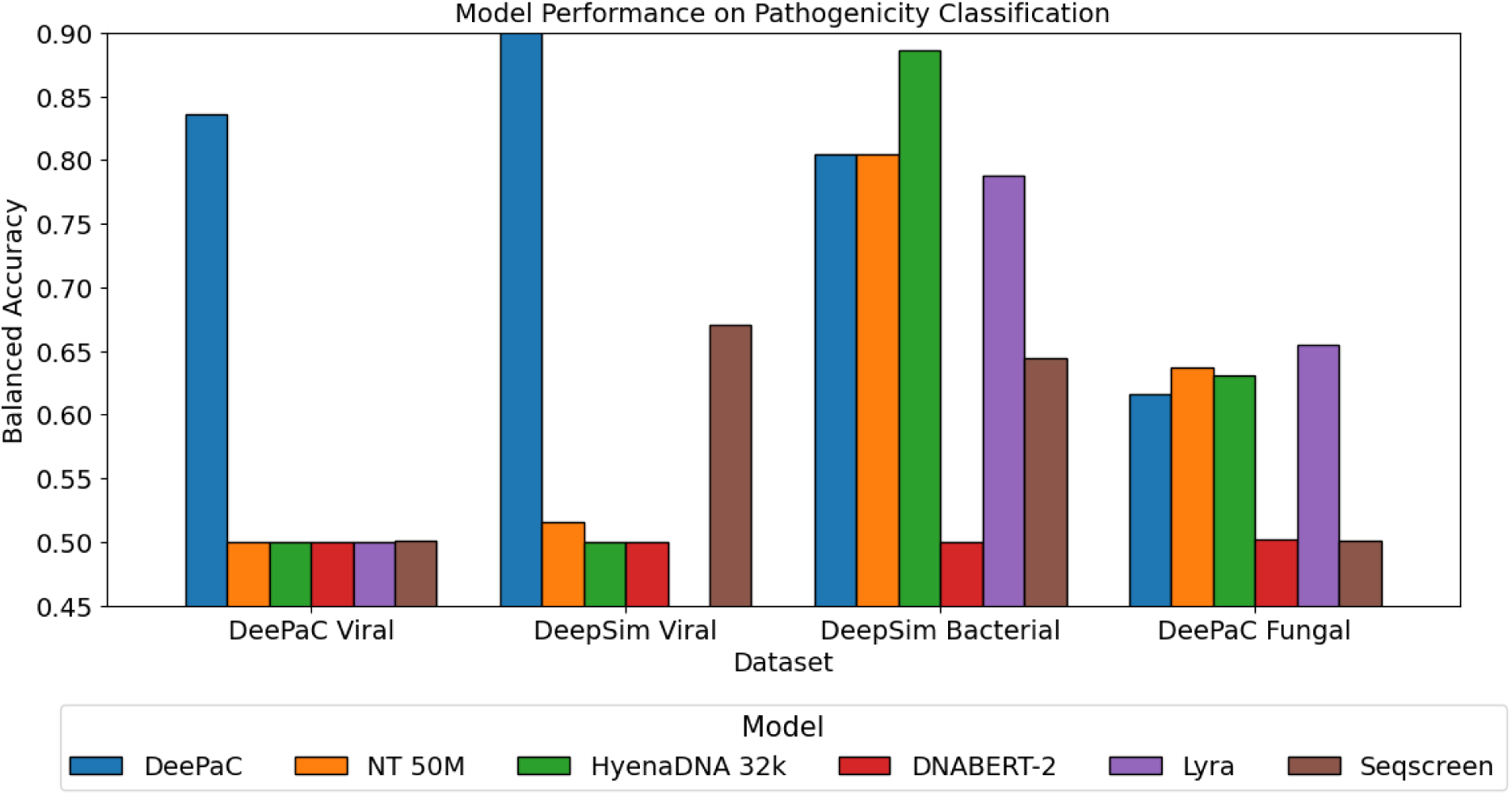
Balanced accuracy of each model across pathogenicity datasets. Performance varied by model and by pathogen type.

Together, these results indicate that pathogenicity signals are less uniformly captured by a single architecture than taxonomic or AMR signals, and instead depend more strongly on organism class and training distribution. Within this framework, DeePaC was most effective for viral pathogenicity, HyenaDNA for bacterial pathogenicity, and Lyra for fungal pathogenicity, supporting a model-matched rather than one-size-fits-all approach to pathogenicity prediction.

### Use case example: Integration of MicrobeRT into TaxTriage as a practical complement to metagenomic workflows

Metagenomic sequencing is a powerful tool enabling untargeted sample analysis. Challenges to its widespread adoption include the complexity and variability of existing analysis tools, as well as the computational requirements to implement them. TaxTriage [5] is an open-source bioinformatics pipeline that mitigates these technical challenges and enables rapid and reproducible processing and analysis of sequence reads. Built using NextFlow [20], the TaxTriage pipeline is comprised of a flexible set of modules that are run sequentially, ultimately outputting an Organism Discovery Report (ODR) as well as a quality control file. The TaxTriage pipeline is already in use at multiple public health laboratories in the United States, and as an open-source pipeline, it is widely available to other public health practitioners, researchers, and laboratory facilities worldwide.

By incorporating our best-performing AI/ML-enabled classification tools into TaxTriage as an optional additional module, we enabled a wide range of potential users to easily leverage the appropriate fine-tuned gLM for taxonomic classification tasks. Users can optionally enable AI/ML taxonomic classification, using the appropriate model for long-read sequences (HyenaDNA-1M) or short-read sequences (NT-50M). Users specify what level of confidence should be included in potential classifications output to the report; for instance, they may choose to report only the most likely taxonomic classification, or they might set the input flag to show the top X most likely classifications, or those with a confidence level *≥* Y %. Taxonomic classification results and associated confidence scores print directly to the ODR alongside results from the standard bioinformatics classification tools.

The integration of MicrobeRT within TaxTriage is shown schematically in Figure 8. Aligned reads were reduced to representative candidate sequences before prediction. Specifically, positional binning and MMseqs2 clustering were used to collapse redundant aligned regions, after which representative sequences were passed to the appropriate MicrobeRT model and predictions were mapped back to the original BAM alignments to supplement the ODR tables. This reduction strategy substantially lowered computational cost. For HyenaDNA, downsampling reduced wall time from 154 to 7.1 min for long reads and from 124.5 to 6.0 min for short reads (Table 1). On the downsampled benchmark sets, NT completed inference in 34.3 min for long reads and 32.5 min for short reads.

**Figure 8:**
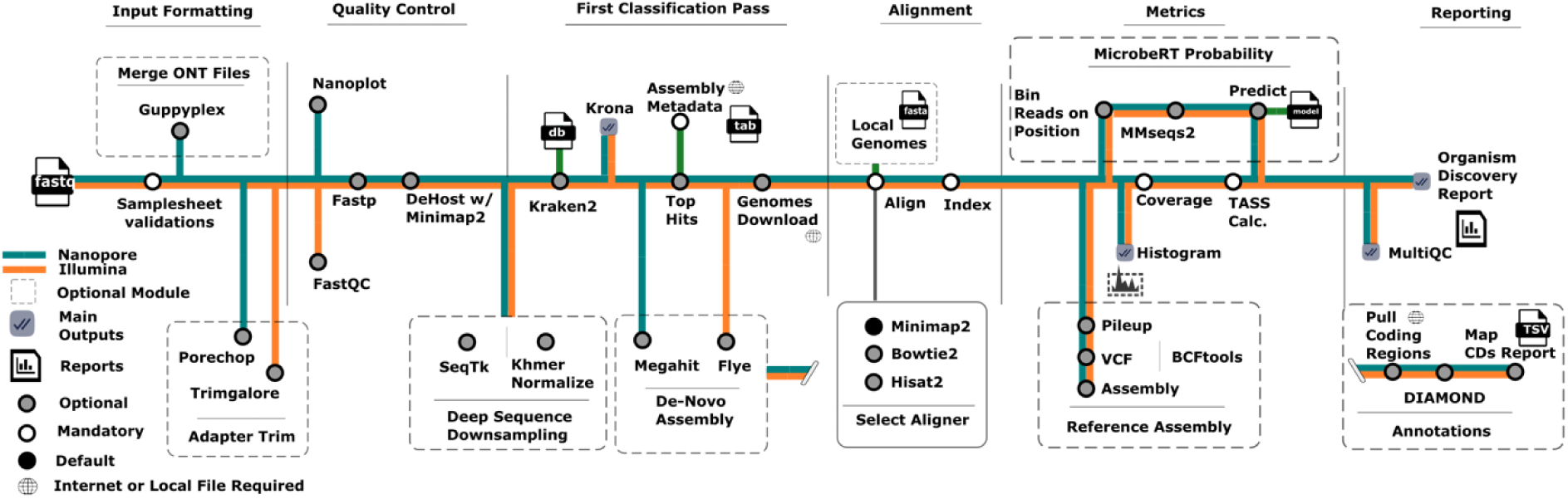
Integration of MicrobeRT (top right) within the TaxTriage workflow.

**Table 1:**
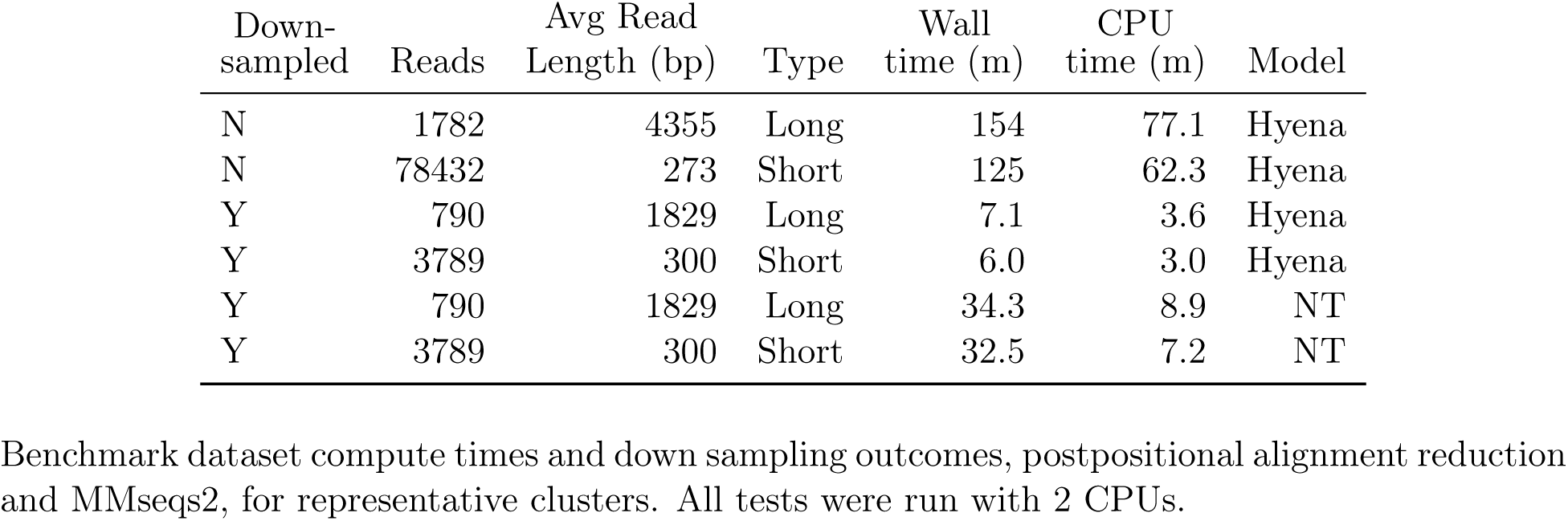
Runtime and dataset characteristics across models and sampling settings.

| Down-sampled | Reads | Avg Read Length (bp) | Type | Wall time (m) | CPU time (m) | Model |
|---|---|---|---|---|---|---|
| N | 1782 | 4355 | Long | 154 | 77.1 | Hyena |
| N | 78432 | 273 | Short | 125 | 62.3 | Hyena |
| Y | 790 | 1829 | Long | 7.1 | 3.6 | Hyena |
| Y | 3789 | 300 | Short | 6.0 | 3.0 | Hyena |
| Y | 790 | 1829 | Long | 34.3 | 8.9 | NT |
| Y | 3789 | 300 | Short | 32.5 | 7.2 | NT |

To demonstrate performance within TaxTriage, a test run with standard configuration and parameters was completed and relevant benchmarks computed (Table 1). The dataset used was the same as the example Samplesheet available for all test runs in TaxTriage, containing 5-10 viruses and bacteria annotated in the pathogen annotation sheet. An example output of the MicrobeRT NT model formatted is shown in Table 2. Both the Hyena and NT probability were taken post-clustering and averaged across all species in the ODR, regardless of whether the row itself is a strain/subspecies (Table 3). In addition, the ODR includes an optional column if the MicrobeRT module is enabled (Figure 9). MicrobeRT outputs were formatted for direct incorporation into TaxTriage reporting, including probability scores mapped to taxonomic identifiers and integrated with existing ODR fields (Figure 9). This design enables users to review MicrobeRT predictions within the same reporting structure used for conventional classifiers rather than through a separate postprocessing workflow.

**Figure 9:**
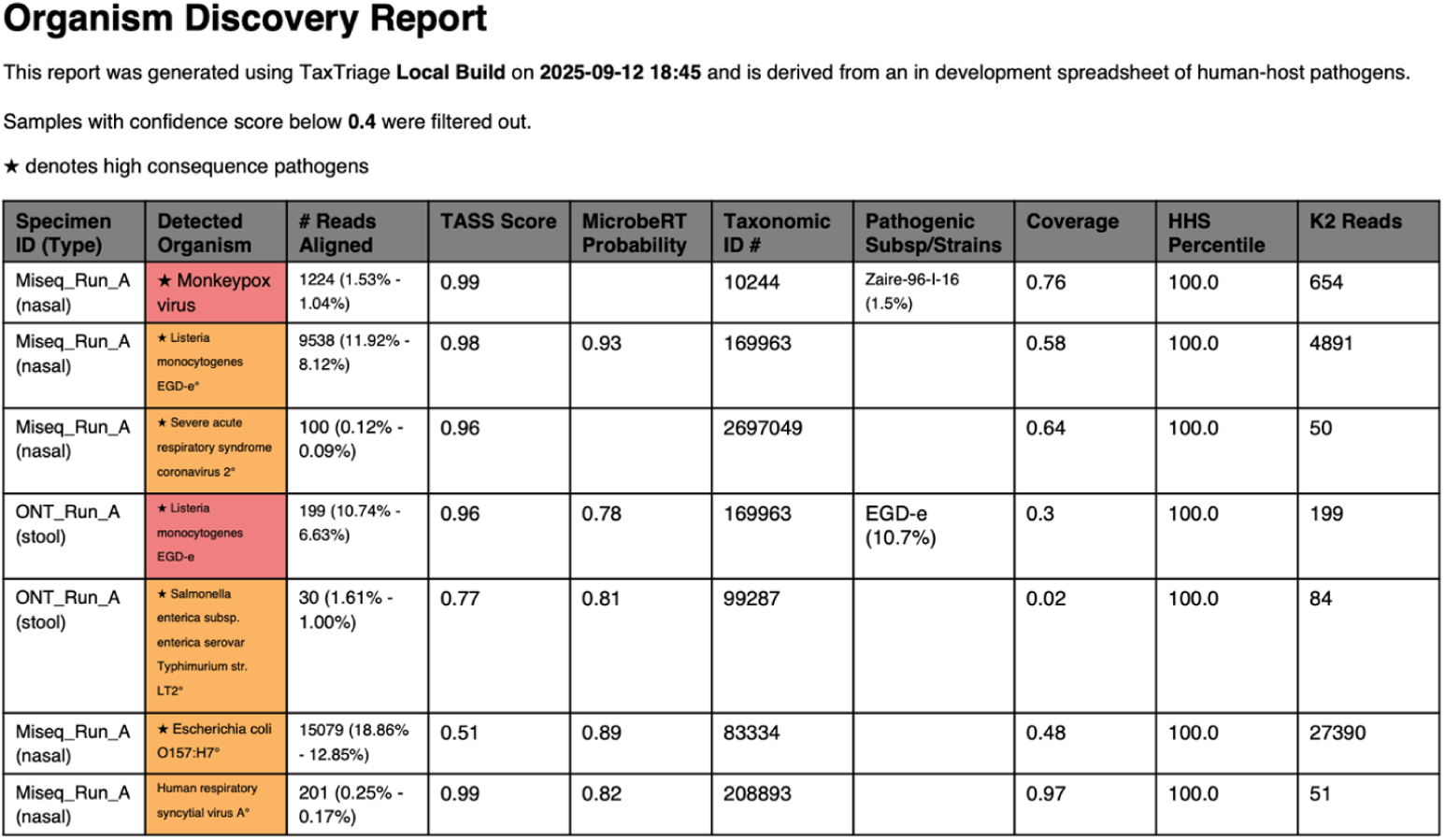
Performance of TaxTriage’s submodules for MicrobeRT probability compared to TASS Score, alignment counts, and K2 Reads classified. The MicrobeRT classifier can be optionally enabled during TaxTriage runs for side-by-side comparison of results. Headers and their respective meanings are as follows: Specimen ID (type): The self-assigned sample identifier for a given set of reads; Detected Organism: The identified organism name; # Reads Aligned: The total number of reads aligned from the alignment step(s) of the workflow; TASS Score: The post-alignment score derived from a novel algorithm in TaxTriage; MicrobeRT Probability: The predicted probability from the MicrobeRT-trained models classification method; Taxonomic ID #: The NCBI taxonomic identifier of the organism; Pathogenic Subsp/Strain: If available, any strains under the species level that have been identified; Coverage: The breadth of coverage of the alignment; HHS Percentile: How relative quantity of the reads aligned compares to the anticipated amount in a given clinical sample of the same type; K2 Reads: Number of Kraken2 reads assigned to the organism at the LCA species-level.

**Table 2:** Example output of the fine-tuned NT model for taxonomic classification. This file can then be uploaded to TaxTriage ODR for interpretation in a reflex response manner.

| Rank | Taxid | Average | Median |
|---|---|---|---|
| superkingdom | 2 | 0.99 | 1.00 |
| phylum | 204428 | 0.99 | 0.99 |
| genus | 45989 | 0.99 | 0.99 |
| genus | 2370 | 0.95 | 0.95 |
| species | 813 | 0.92 | 0.92 |
| species | 2371 | 0.91 | 0.91 |
| family | 1706371 | 0.91 | 0.91 |
| phylum | 1224 | 0.90 | 0.98 |
| phylum | 1297 | 0.86 | 0.86 |
| genus | 638847 | 0.84 | 0.84 |
| family | 27974 | 0.83 | 0.83 |
| phylum | 1853220 | 0.83 | 0.83 |
| species | 2527972 | 0.81 | 0.81 |
| species | 521 | 0.80 | 0.80 |
| superkingdom | 2759 | 0.79 | 0.85 |
| family | 2691359 | 0.78 | 0.78 |
| genus | 2147 | 0.78 | 0.78 |
| family | 1903416 | 0.74 | 0.74 |
| genus | 133193 | 0.74 | 0.74 |
| family | 1033997 | 0.74 | 0.74 |
| phylum | 203682 | 0.73 | 0.76 |

**Table 3:**
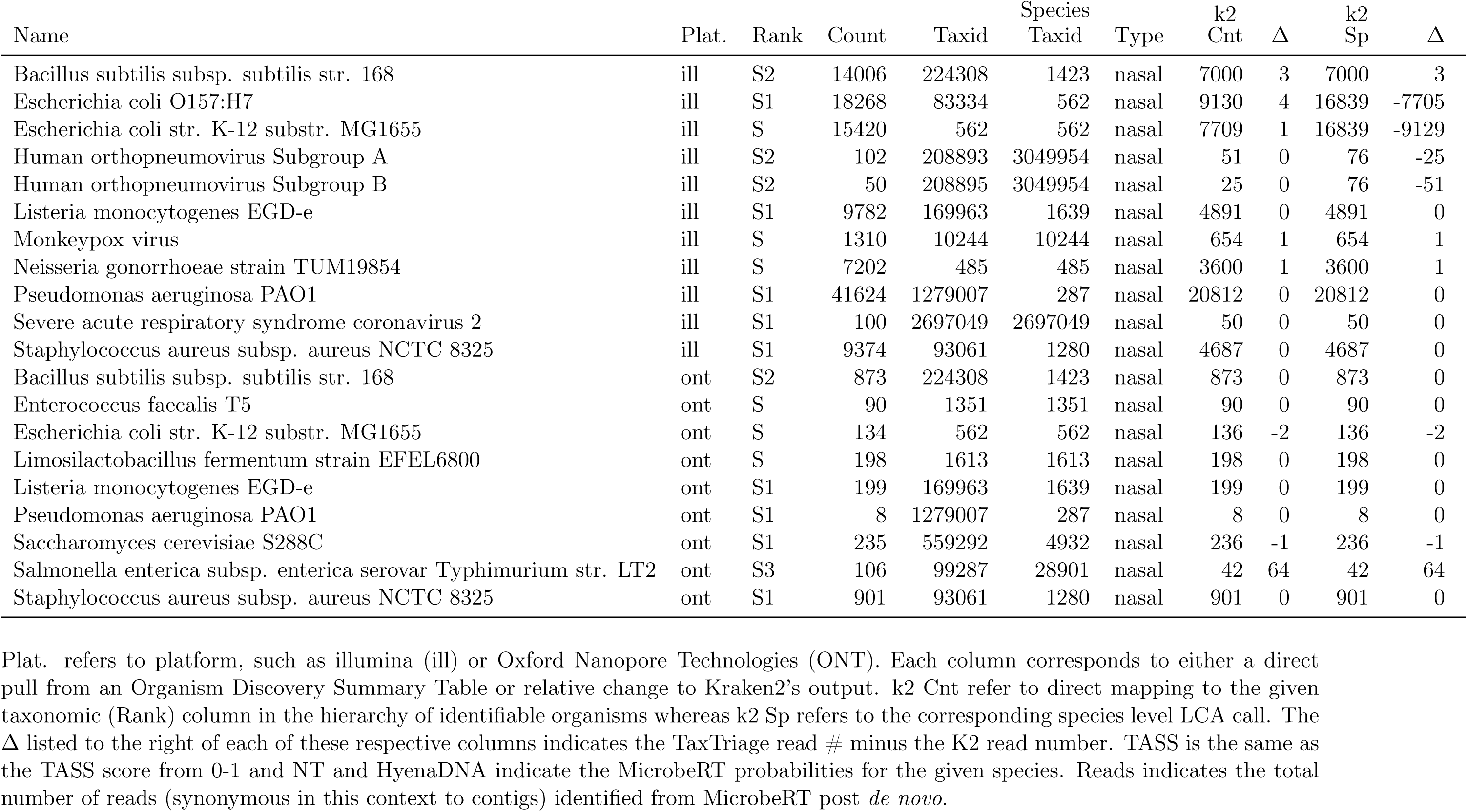
Example output of the test run of each model within TaxTriage.

|  |  |  |  |  | Species |  | k2 |  | k2 |  |
|---|---|---|---|---|---|---|---|---|---|---|
| Name | Plat. | Rank | Count | Taxid | Taxid | Type | Cnt | Δ | Sp | Δ |
| Bacillus subtilis subsp. subtilis str. 168 | ill | S2 | 14006 | 224308 | 1423 | nasal | 7000 | 3 | 7000 | 3 |
| Escherichia coli O157:H7 | ill | S1 | 18268 | 83334 | 562 | nasal | 9130 | 4 | 16839 | -7705 |
| Escherichia coli str. K-12 substr. MG1655 | ill | S | 15420 | 562 | 562 | nasal | 7709 | 1 | 16839 | -9129 |
| Human orthopneumovirus Subgroup A | ill | S2 | 102 | 208893 | 3049954 | nasal | 51 | 0 | 76 | -25 |
| Human orthopneumovirus Subgroup B | ill | S2 | 50 | 208895 | 3049954 | nasal | 25 | 0 | 76 | -51 |
| Listeria monocytogenes EGD-e | ill | S1 | 9782 | 169963 | 1639 | nasal | 4891 | 0 | 4891 | 0 |
| Monkeypox virus | ill | S | 1310 | 10244 | 10244 | nasal | 654 | 1 | 654 | 1 |
| Neisseria gonorrhoeae strain TUM19854 | ill | S | 7202 | 485 | 485 | nasal | 3600 | 1 | 3600 | 1 |
| Pseudomonas aeruginosa PAO1 | ill | S1 | 41624 | 1279007 | 287 | nasal | 20812 | 0 | 20812 | 0 |
| Severe acute respiratory syndrome coronavirus 2 | ill | S1 | 100 | 2697049 | 2697049 | nasal | 50 | 0 | 50 | 0 |
| Staphylococcus aureus subsp. aureus NCTC 8325 | ill | S1 | 9374 | 93061 | 1280 | nasal | 4687 | 0 | 4687 | 0 |
| Bacillus subtilis subsp. subtilis str. 168 | ont | S2 | 873 | 224308 | 1423 | nasal | 873 | 0 | 873 | 0 |
| Enterococcus faecalis T5 | ont | S | 90 | 1351 | 1351 | nasal | 90 | 0 | 90 | 0 |
| Escherichia coli str. K-12 substr. MG1655 | ont | S | 134 | 562 | 562 | nasal | 136 | -2 | 136 | -2 |
| Limosilactobacillus fermentum strain EFEL6800 | ont | S | 198 | 1613 | 1613 | nasal | 198 | 0 | 198 | 0 |
| Listeria monocytogenes EGD-e | ont | S1 | 199 | 169963 | 1639 | nasal | 199 | 0 | 199 | 0 |
| Pseudomonas aeruginosa PAO1 | ont | S1 | 8 | 1279007 | 287 | nasal | 8 | 0 | 8 | 0 |
| Saccharomyces cerevisiae S288C | ont | S1 | 235 | 559292 | 4932 | nasal | 236 | -1 | 236 | -1 |
| Salmonella enterica subsp. enterica serovar Typhimurium str. LT2 | ont | S3 | 106 | 99287 | 28901 | nasal | 42 | 64 | 42 | 64 |
| Staphylococcus aureus subsp. aureus NCTC 8325 | ont | S1 | 901 | 93061 | 1280 | nasal | 901 | 0 | 901 | 0 |

Plat. refers to platform, such as illumina (ill) or Oxford Nanopore Technologies (ONT). Each column corresponds to either a direct pull from an Organism Discovery Summary Table or relative change to Kraken2’s output. k2 Cnt refer to direct mapping to the given taxonomic (Rank) column in the hierarchy of identifiable organisms whereas k2 Sp refers to the corresponding species level LCA call. The Δ listed to the right of each of these respective columns indicates the TaxTriage read # minus the K2 read number. TASS is the same as the TASS score from 0-1 and NT and HyenaDNA indicate the MicrobeRT probabilities for the given species. Reads indicates the total number of reads (synonymous in this context to contigs) identified from MicrobeRT post *de novo*.

Taken together, these results show that MicrobeRT can be deployed as a practical adjunct to existing metagenomic pipelines, improving workflow compatibility and interpretability while keeping sequence-based classification computationally manageable.

## Discussion

Accurate and reproducible pathogen detection remains a central challenge in metagenomic-based diagnostics because current workflows often depend on complex tools, mutable reference databases, and substantial computational resources. Here, we introduce MicrobeRT, an integrated framework and suite of three complementary genomic language models—Nucleotide Transformer (NT), DNABERT, and HyenaDNA—fine-tuned for taxonomic classification, AMR gene detection, and pathogenicity prediction. Across these tasks, MicrobeRT was most effective when used as a complement to, rather than a replacement for, reference-based classifiers and other ML-based approaches. Its greatest advantage is in instances where contextual sequence modeling is likely to matter most, including long-read taxonomic classification, AMR prediction under truncated or flanking-sequence perturbations, and detection of organisms or genes that were well represented during fine-tuning. Together, these findings support the broader premise that pre-trained genomic language models can extend metagenomic analysis beyond exact sequence matching while remaining compatible with established bioinformatics workflows. However, they also reiterate the importance of classical bioinformatic tools and the ongoing need to improve and refine these algorithms.

For taxonomic classification, model performance depended on read length and representation in the fine-tuning data. HyenaDNA performed best on long fragments, whereas NT and DNABERT-2 were more successful in classifying short reads, consistent with their differing context capacities and tokenization schemes. At the same time, Kraken2 retained high performance for sparsely represented taxa, indicating that genomic language models such as those in the MicrobeRT suite do not yet surpass mature reference-based classifiers for broad taxonomic coverage. Thus, the strongest use case for MicrobeRT is as a part of hybrid workflows, in which sequence-based models provide complementary sensitivity for long reads and for taxa that are well supported during fine-tuning, including clinically relevant organisms such as *Bordetella pertussis*.

The AMR results demonstrate that gLMs can learn patterns beyond exact sequence identity; in this case, resistance-associated information. NT and HyenaDNA outperformed DeepARG in binary AMR-associated sequence detection, whereas NT-250M and DNABERT-S outperformed DeepARG in multi-label prediction across full-length, truncated, and flanked sequence inputs. This robustness is relevant for metagenomic applications, in which AMR genes are often recovered as incomplete fragments or embedded within heterogeneous genomic context. Furthermore, accurate AMR gene detection can help inform antibiotic selection and assist in downstream confirmatory testing. Together, these findings indicate that genomic language models can contribute clinically relevant AMR information even when reference similarity alone is insufficient.

In contrast to taxonomy and AMR detection, pathogenicity prediction showed stronger dataset- and kingdom-specific effects. DeePaC performed best on viral datasets, HyenaDNA performed best on the bacterial benchmark, and Lyra achieved the highest performance on the fungal dataset. This heterogeneity suggests that pathogenicity signals are less uniformly captured by a single model family and may depend more strongly on organism class, sequence characteristics, and training distribution. These results support a modular framework in which different architectures can be selected according to the biological regime of interest.

The MicrobeRT framework, including code and fine-tuned models, is available open-source, enabling rapid and reproducible assessment of the performance of newly published models alongside existing fine-tuned models and SOTA classifiers. Importantly, MicrobeRT is designed to be modelagnostic and can incorporate externally developed models. Additional models such as the ICCTax model [21], which fine-tunes HyenaDNA for taxonomic classification with a focus on marine and wastewater datasets, and ViTax [22], which is specifically developed for viral classification, are distinct from the models introduced in this work but can be readily integrated into the MicrobeRT framework and evaluated within a consistent benchmarking pipeline.

Another practical strength of this study is that MicrobeRT was evaluated not only as a standalone modeling framework but also as an optional module within TaxTriage. This integration demonstrates that gLMs can be incorporated into established metagenomic pipelines without displacing familiar tools and reporting formats, thereby lowering barriers to adoption in public health and research settings.

Since the evaluated architectures differ substantially in memory requirements and context windows, identical training and input regimes were not feasible across all models and tasks, reducing conformity in testing. Additionally, the performance of MicrobeRT models was related to representation in the fine-tuning data, indicating that data with broader taxa and resistance-class coverage are needed for improved fine-tuning for these three tasks. Furthermore, the rapidly expanding gLM landscape suggests that newer models and architectures may further improve performance. In future work, we hope to additionally leverage powerful and recently published models such as GenomeOcean [23] and METAGENE-1 [24].

MicrobeRT demonstrates the feasibility of an approach that uses fine-tuning pre-trained genomic language models to perform key classification tasks to improve and enhance metagenomic sequence analysis. By evaluating gLM results alongside existing SOTA bioinformatics classification tools, the relative strengths and weaknesses of each approach were demonstrated in a reproducible and comprehensive pipeline. Integration of the taxonomic classification tools with TaxTriage will enable end users to easily access and run the relevant models within an established bioinformatics workflow. MicrobeRT has been developed open-source, enabling future development to include additional model families, datasets, and fine-tuning approaches. The MicrobeRT tools serve as a starting framework for extending metagenomic analysis beyond exact sequence matching while preserving compatibility with current bioinformatics workflows.

## Materials and Methods

### Taxonomy data

Representative genomes were downloaded from NCBI RefSeq and GenBank using ncbi-genome-download [25]. RefSeq provided representative genomes for archaea, bacteria, and viruses and Gen-Bank expanded coverage with complete and chromosome-level bacterial assemblies, viral assemblies across all completeness levels, and fungal assemblies (complete, chromosome, and scaffold). Representative host genomes (human, cow, dog, cat, pig, wheat, corn) and AMR reference sequences were also incorporated. Genomes were randomly fragmented into 750–3,500 bp subsequences, then clustered with MMSeqs2 at 90% and 70% identity to remove redundancy, producing *∼*155.7M nonredundant sequences. A *∼*1% hold-out set (1.57M sequences, 500–1,750 bp) was reserved for testing. Additional details can be found in the supplementary information (See SI: Dataset Collection, Cleaning and Labeling - Taxonomy).

### Antimicrobial resistance data

AMR-positive sequences were identified using the Resistance Gene Identifier (RGI) under Perfect (exact matches to CARD) and Strict (novel variants with key mutations) paradigms [26]. Together with negatives extracted from the taxonomy database, there was a total of *∼*672k positives and 3.6B negatives. Sequences were fragmented into 140–475 bp subsequences, then clustered (100% identity for positives, 70% for negatives), yielding 300,091 unique positive fragments and 86.9M negatives. To assess model performance under different sequence length conditions, five test sets were obtained: full-length AMR genes, truncated genes (140–450 bp), and genes flanked with 50 bp, 100 bp, or 50–2,500 bp random context. AMR-positive sequences were further labeled by drug class using DeepARG [13] categories, with 28 of 30 classes represented and negative controls generated using positional information to extract sequence fragments that did not map to AMR genes. Additional details can be found in the SI Appendix.

### Pathogenicity data

Fungal genomes were sourced from a curated dataset of pathogenic and non-pathogenic fungi genomes [27], totaling 42.6M sequences after fragmentation into 500–1,750 bp subsequences. To expedite training time and limit redundancy, these were then down selected to 20% of the total sequences. Oxford Nanopore bacterial and viral reads were simulated using DeepSimulator version 1.5 [28, 29], with 85% fidelity, due to the error rate specified in the software simulation. Pathogenicity labels were assigned using CDC bioterrorism categories [30] and NIH biosafety risk groups [31]. Redundancy was further reduced with BBNorm [32] and MMSeqs2 [33] clustering, producing 109M labeled sequences (63.5M bacterial, 2.9M viral, 42.6M fungal)(30 GB positive and 28 GB negative). A second viral dataset, DeePaC Viral [34], was tested for comparison to performance on the simulated viral dataset, DeepSim Viral. Additional details can be found in the SI Appendix.

### Model Fine-tuning

The models fine-tuned in this work were all built using the following pre-trained genomic foundation models: Nucleotide Transformer (NT), DNABERT-2 and DNABERT-S, and HyenaDNA. These models differ in architecture, pre-training data, and tokenization.

- **Nucleotide Transformer (NT) [1]:** Transformer models ranging from 50M to 2.5B parameters, pre-trained on 174B nucleotides from 850+ species. Uses non-overlapping 6-mer tokenization with a maximum sequence length of 1,000 bp.
- **DNABERT-2 [2]:** A 117M parameter transformer adapted from BERT13, pre-trained on 32.49B nucleotides from 135 species. Uses Byte Pair Encoding (BPE) tokenization to produce variable-length tokens, better capturing genome structure than fixed k-mers (sliding windows of short subsequences). Pre-trained with a maximum sequence length of 512 bp. We also considered DNABERT-S [3], which is built off DNABERT-2 and further optimized through supervised training for species-level classification tasks.
- **HyenaDNA [4]:** Family of non-transformer models (maximum 6.6M parameters) pre-trained on the human reference genome. Unlike encoder-only transformers, it is decoder-only and replaces attention with the Hyena operator, enabling efficient single-nucleotide processing and context lengths up to 1 million tokens. Excels at long-read sequencing data.

HyenaDNA models were fine-tuned on long fragments ranging from 500 to 1,750 bps. Finetuning was done on up to eight NVIDIA H100 GPUs for five epochs with a weight decay of 0.01, a cosine learning rate scheduler (0.05 warmup ratio), and bf16 precision. Early stopping with a patience of three epochs was implemented to prevent overfitting. The per-device batch size was set to the maximum permitted by GPU memory. To handle the extremely large fine-tuning dataset size, streaming-based batched tokenization and training were employed. Exploratory experiments on 0.05% data subsets were used to select optimum learning rates. We also evaluated curriculum-style training across taxonomic ranks, but observed no performance gains compared to direct multi-rank fine-tuning. In prior work, we additionally conducted extensive evaluation of parameter-efficient fine-tuning (PEFT) approaches, including LoRA and (IA)^3^ [35], but observed that they were ineffective for the downstream task of taxonomic classification across multiple detailed experiments. Additional details can be found in the SI Appendix.

The same training strategy was applied for fine-tuning the NT and DNABERT models, with the exception that training was performed on a further processed version of the taxonomic dataset. Because transformer model memory scales quadratically with sequence length, training on the original dataset (up to 1,750 bp) would have required over a month, making it impractical. To mitigate this, sequences were further chunked to within 300 bp, which increased the overall dataset to 614,410,126 sequences but reduced the memory footprint per batch, allowing training to finish reasonably within two weeks. A hold-out set of 606,971 sequences was reserved for validation.

## Acknowledgments and funding sources

This work was supported by funding from the U.S. Centers for Disease Control and Prevention through the Office of Readiness and Response under Contract # 75D30124C20202. The authors acknowledge funding from the Johns Hopkins University Applied Physics Laboratory Independent Research and Development Program. We would like to thank Dr. Matt Mauldin, Dr. Zahra Chaudhry, Dr. Paul Nicholas, Dr. Philip Graff, and Dr. Sarah Grady for their valuable insights and feedback during the completion of this work.

## Author contributions

J.R. and V.N. designed the research; J.R. and M.G. supervised the research; V.N., D.B., and D.J. developed the model fine-tuning and evaluation pipeline; S.T., B.M., and S.C. curated and processed the sequence data; and M.G., S.C., B.M., V.N., D.J., S.T., and D.B. wrote the paper.

