## Supplementary Information for "MicrobeRT: A Framework and Suite of Fine-tuned Language Models for Genomic Sequences"

#### Dataset Collection, Cleaning and Labeling

Here we describe the data collection methodology for the three downstream classification tasks: taxonomy, antimicrobial resistance, and pathogenicity. For all three tasks, datasets were split into training and test sets with class balance maintained. Additionally, for the pathogenicity task, the data were further stratified into viral, bacterial, and fungal reads.

##### Taxonomy

For taxonomic classification, training data were obtained using ncbi-genome-download [1]. Genomes were retrieved from the NCBI RefSeq [2] and GenBank [3] databases according to specified options. From RefSeq, representative genomes were downloaded for bacteria, viruses, and archaea (Table 1). To expand coverage, we included all complete and chromosome-level bacterial assemblies from GenBank, as well as viral assemblies across all levels. For fungi, complete, chromosome, and scaffold assemblies were downloaded from GenBank. For comparison, a single representative assembly was retrieved for potential host organisms, including human, cow, dog, domestic cat, pig, wheat, and corn. Additionally, sequences from the NCBI Antimicrobial Resistance (AMR) Reference Gene Catalog [4] were downloaded, mapped to their taxonomy, and incorporated into the taxonomy database. In total, the entire uncompressed training data is 258GB in size. To minimize redundancy, a cluster-based down-selection was applied. Full-length sequences were first randomly fragmented into subsequences of 750-3500 base pairs (bp). These subsequences were clustered using MMSeqs2 at 90% identity within each taxonomic family, and one representative sequence was retained per cluster. The resulting representatives were then clustered again at 70% identity, with a single sequence selected from each cluster to form the final dataset. In addition, derivative datasets of defined sequence lengths were generated to benchmark trained genomic tokenizers. A hold-out set of 1% of the total dataset, comprising 1,572,987 sequences, was reserved for testing. Sequence lengths ranged from 500–1750 bp.

##### Antimicrobial Resistance

To obtain AMR sequences for training, the Resistance Gene Identifier (RGI) was used [5]. RGI predicts antibiotic resistomes from protein or nucleotide data based on homology and single nucleotide polymorphism (SNP) models. RGI first predicts complete open reading frames (ORFs) using Prodigal, then analyzes the resulting protein sequences under user-specified paradigms: Perfect, requiring

Table 1: Summary of sequence representation of taxa present in the database for taxonomic prediction.

| Superkingdom / Kingdom | Representation (n) |
|---|---|
| Archaea | 1,252,736 |
| Bacteria | 145,122,383 |
| Eukaryota | 8,633,180 |
| Viruses | 717,342 |
| Total | 155,725,641 |

Table 2: Summary of sequence representation present in the training data for AMR prediction (binary).

| AMR Status | Representation (n) |
|---|---|
| Negative | 86,866,041 |
| Positive | 3,000,091 |
| Total | 89,866,132 |

exact matches to curated CARD references, or Strict, which detects novel variants of known AMR genes and screens for key mutations. To substantially expand the training dataset, both the Perfect and Strict algorithms were applied across the taxonomy database. Combined, these datasets contained a total of 672,392 AMR-positive sequences and 3,598,448,873 AMR-negative sequences. These sequences were then randomly fragmented into lengths ranging from 140–475 bp, as AMR-positive fragments were, at minimum, 150 bp. MMSeqs2-based clustering [6] was then applied to reduce redundancy where AMR-positive fragments were clustered at 100% sequence identity, and AMR-negative fragments were clustered at 70% sequence identity, and one representative sequence was retained per cluster. As seen in Table 2, this resulted in 3,000,091 unique AMR-positive fragments and 86,866,041 unique AMR-negative sequences.

To assess model performance under different sequence length conditions, five test sets were generated. The first contained full-length AMR sequences from the CARD and NCBI AMR Reference databases. The second included partial AMR sequences randomly truncated to 140–450 bp. The remaining three consisted of full AMR sequences flanked on both ends by 50 bp, 100 bp, or randomly distributed lengths of 50–2500 bp. Table 3 lists the number of sequences in each set.

In addition to a binary classification of sequences resistance, information about the specific antibiotic to which each sequence shows resistance was extracted from the RGI output to assess the models' ability to perform multi-class classification. To generate negative controls, positional information was used to extract sequence fragments that did not map to antimicrobial resistance

Table 3: Summary of test sets and number of sequences in each for AMR classification.

| Test Set | Representation (n) |
|---|---|
| Whole AMR | 14,069 |
| Partial AMR | 20,000 |
| 50 bp flank | 12,093 |
| 100 bp flank | 12,093 |
| Random flanks between 50 and 2500 bp | 12,093 |

genes (ARGs).

The extracted antibiotic drug classes for the 300,091 positive training sequences were mapped to 30 labels, as defined by DeepARG [7]. As seen in Table 4, of the 30 DeepARG labels, 28 were represented in the training dataset. Similarly, for each of the test datasets, each sequence was mapped to the 28 out of 30 DeepARG labels present in the training data. Not all sequences from the pulled test datasets were retained for minulti-class assessment as some of the labels in the test data were not present in the training data. The totals in the tables reflect the number of sequences used in the training and test data for multi-class prediction.

#### Pathogenicity

Bacterial, viral, and fungal data were obtained for pathogenicity classification (Table 5). Fungal pathogens were sourced from a curated dataset of pathogenic and non-pathogenic fungi genomes published in 2022 [8]. Accessions from this dataset were used to download 24,650,287 genomes ( 40GB). Of these, 5,769,444 sequences were pathogenic and 18,880,842 were non-pathogenic. Sequences were then fragmented into random lengths of 500–1750 bp for a total of 42,664,025 sequences in the original training set. To expedite training time and limit redundancy, these were then down selected to 20% of the total sequences. A dataset previously curated by simulating Oxford Nanopore Long reads with DeepSimulator (DeepSim) version 1.5 [9, 10] was used for the Bacterial and Viral sequences. These are usually a few kb in length and have relatively low quality ( 85% fidelity) due to the introduction of simulated error by the software. Human pathogenicity was determined from the U.S. Centers for Disease Control and Prevention (CDC) bioterrorism categories [11] and the National Institutes of Health (NIH) biosafety risk groups [12]. The total size of the dataset was 1.4 TB.

Because of training constraints, the total dataset size was reduced using the following tools designed to eliminate redundant sequences:

1. BBMap's BBNorm [13] submodule to Downsample
   - Positive: 767 GB reduced to 39 GB (18 kmer length)
   - Negative: 659 GB reduced to 29 GB (14 kmer length, 18 was >200 GB)
2. MMSeqs2 Clustering [6]
   - Bin each pos/negative Downsample file on phylum (one FASTA per phylum of all taxa)
   - Generates a FASTA file
   - Run MMSeqs2 linclust on each FASTA file
   - Pull ONLY representative sequences from each cluster
   - Final Sizes:
     - Positive: 39 GB reduced to 30 GB
     - Negative: 29 GB reduced to 28 GB

The relative classes were imbalanced according to phylum as indicted by the NIH Risk Levels and CDC Priority Pathogen lists (Figure 1). Additionally, the breakdown of Superkingdom across the simulated datasets were: Bacteria at 4,146,891 and 4,635,983 for negative and positive, respectively, while for viral reads, the counts were much smaller at 318,033 and 1,037,789 respectively (Figure 2).

Table 4: Summary of sequence representation present in the training and testing data for AMR prediction (multi-class).

| Antibiotic Drug Class | Training (n) | Whole AMR (n) | Partial AMR (n) | 50 bp flank (n) | 100 bp flank (n) | Random flanks (50–2500 bp) (n) |
|---|---|---|---|---|---|---|
| acriflavine | 580 | | | | | |
| aminocoumarin | 16,217 | 297 | 1 | 1 | 1 | 1 |
| aminoglycoside | 13,062 | | 383 | 182 | 182 | 182 |
| bacitracin | 1,128 | | | | | |
| beta-lactam | 10,299 | 5,809 | 2,850 | 4,906 | 4,906 | 4,906 |
| bleomycin | 6 | | | | | |
| fosfomycin | 3,658 | 11 | 1,229 | 37 | 37 | 37 |
| fusidic-acid | 51 | 7 | 10 | 9 | 9 | 9 |
| glycopeptide | 65,297 | | | | | |
| kasugamycin | 18 | | | | | |
| linezolid | 11 | | | | | |
| macrolide-lincosamide-streptogramin | 4,994 | 164 | 422 | 131 | 131 | 131 |
| multidrug | 9,359 | 4 | 10 | 7 | 7 | 7 |
| mupirocin | 82 | | 8 | 3 | 3 | 3 |
| peptide | 686 | 109 | 730 | 210 | 210 | 210 |
| phenicol | 2,466 | 84 | 1,443 | 59 | 59 | 59 |
| pleuromutilin | 1,271 | 12 | 1,271 | 28 | 28 | 28 |
| polymyxin | 13,479 | | | | | |
| puromycin | 3,619 | 19 | 32 | 23 | 23 | 23 |
| quinolone | 22,297 | 183 | 66 | 162 | 162 | 162 |
| rifampin | 8,648 | 29 | 41 | 43 | 43 | 43 |
| roxithromycin | 508 | | | | | |
| streptothricin | 72 | | | | | |
| sulfonamide | 191 | 7 | 750 | 6 | 6 | 6 |
| tetracycline | 100,124 | 81 | 396 | 109 | 109 | 109 |
| triclosan | 13,045 | | | | | |
| trimethoprim | 8,801 | 120 | 111 | 102 | 102 | 102 |
| tunicamycin | 110 | | | | | |
| viomycin | 12 | | | | | |
| **Total** | 300,091 | 6,936 | 9,753 | 6,018 | 6,018 | 6,018 |

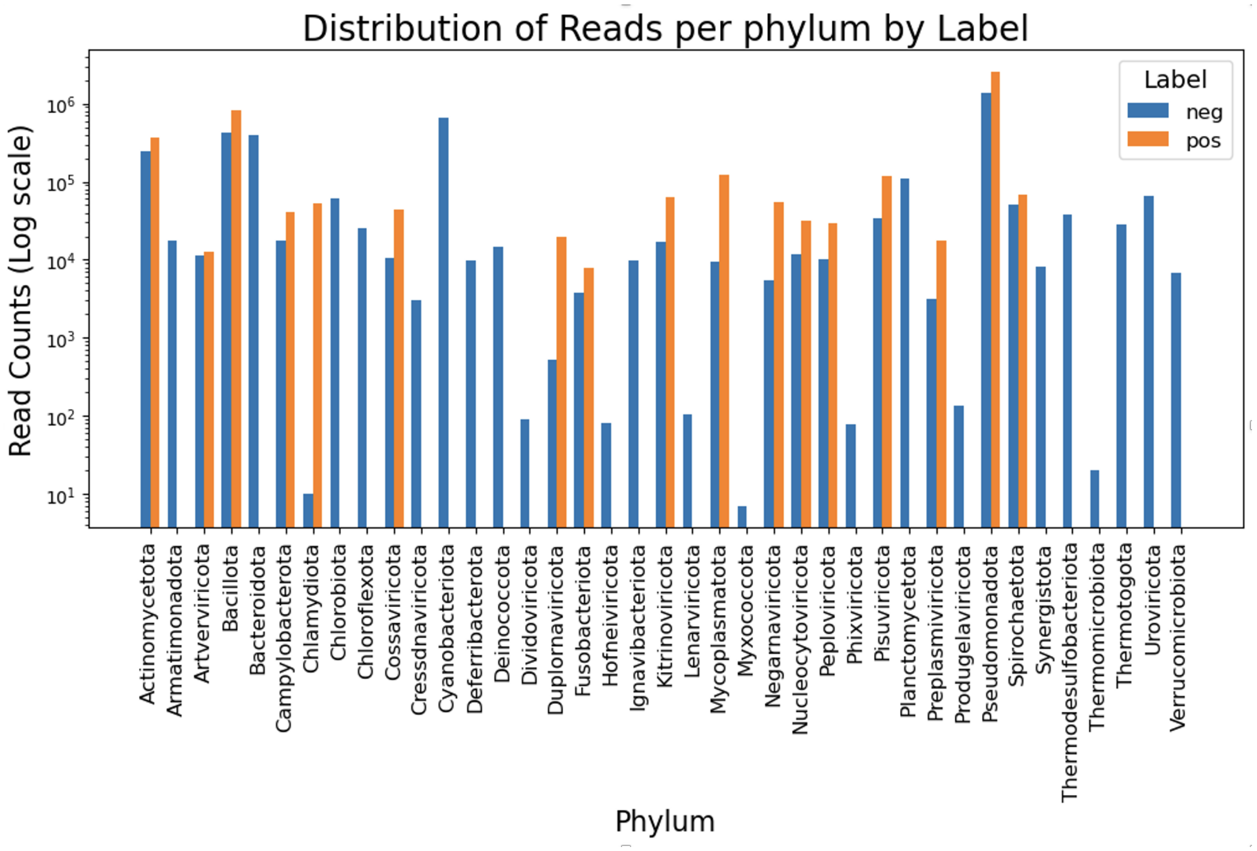


Figure 1: Final distribution of Bacterial and Viral read counts by label and phylum.

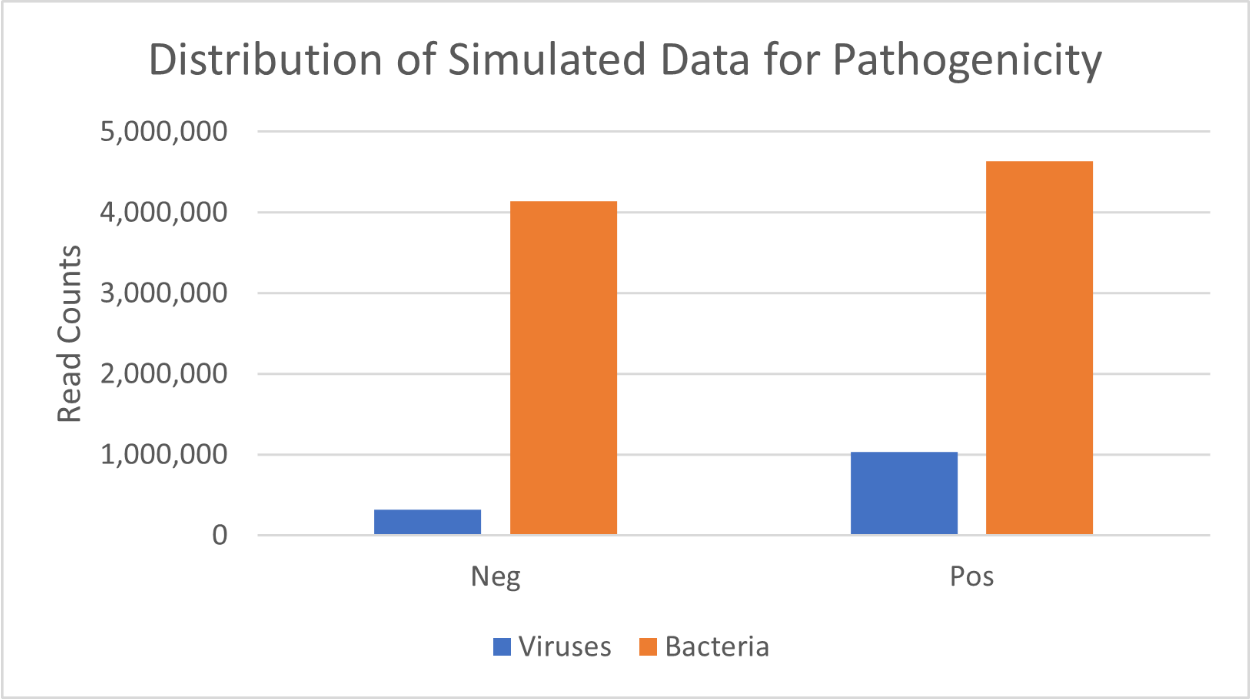


Figure 2: Distribution of the simulated viral and bacterial datasets of 1.4 TB using deepsimulator [9] and then down sampled using MMseqs2 and BBNorm.

Table 5: Summary of sequence representation in the training data for pathogenicity identification

| Pathogenicity Kingdom | Representation (n) |
|---|---|
| Bacteria | 63,482,834 |
| Viral | 2,898,045 |
| Fungal | 42,664,025 |
| Total | 109,044,904 |

### Model Training

#### Model Architecture

The models fine-tuned in this work were all built using the following pre-trained genomic foundation models: Nucleotide Transformer (NT), DNABERT-2 and DNABERT-S, and HyenaDNA. These models differ in architecture, training data, and tokenization. The fine-tuning and evaluation pipeline developed in this work is available on GitHub (https://github.com/jhuapl-bio/microbert), and all fine-tuned models produced in this study are available on Hugging Face (https://huggingface.co/jhuapl-bio/microbert).

For each classification task, an additional classification head was appended to the base architecture for each label being classified. Each head was implemented as a linear layer mapping the hidden dimension of the base model to the number of classes for that label. In the case of taxonomic classification, each of the classification heads is independent of each other. The implementation of a fully hierarchical architecture, where outputs from one rank inform the next, is left for future exploration. During a forward pass, input token ids and attention masks are processed by the base model to produce token-level hidden states. We extract the last hidden state and apply mean pooling across the tokenized sequence to obtain a fixed-size embedding for each input (e.g. 512 dimensional embeddings for NT-50M and NT-100M). To compute mean pooling, we expand the attention mask to match the dimensions of the hidden states, then compute the sum of the token embeddings weighted by the attention mask, and divide by the total number of valid (non-padded) tokens. This ensures only meaningful tokens contribute to the final sequence embedding. Finally, the mean pooled embeddings are passed through each of the classification heads to produce logits corresponding to each label. During training, a separate cross-entropy loss is computed for each label, and the overall loss is calculated as the sum of these individual losses. Our fine-tuning pipeline leverages an iterable dataset to stream tokenized sequences during training, avoiding the need to load the entire dataset into memory and thereby preventing out-of-memory errors. Rather than pre-computing and storing all inputs, sequences are processed on-the-fly, enabling efficient handling of large-scale genomic datasets. To further support distributed training, the dataset is sharded across devices to operate on distinct subsets of the data in parallel. At each epoch, the validation F1 macro score [14] was tracked in order to save off the checkpoint of the corresponding model. The whole design of the script considers memory and GPU safety for large biological sequence datasets.

Beyond the primary pre-trained foundation models explored, we also evaluated a recent subquadratic state space architecture, Lyra [15]. Unlike the fine-tuned models, which leverage representations learned during large-scale genomic pre-training, Lyra is trained more directly for the task yet still demonstrates competitive generalization performance. The model first encodes input sequences and then processes them through two PGC blocks (gated convolutions with RMSNorm) and one S4D block (a structured state space model with a diagonal kernel). It incorporates standard deep learning components, including residual connections, dropout, and mean pooling across the sequence length. This design balances local feature extraction through convolutional layers (PGC) with the ability to capture long-range dependencies via the S4D block. Since the authors of [15] only provide a notebook demo script, a separate pipeline outside of the existing fine-tuning approach was created to handle evaluation on our large genomic datasets. Changes were incorporated to the training logic to make it more consistent with the best practices used throughout the previous benchmarking and evaluations. The first of these refinements was the data handling. Our MicrobeRT fine-tuning process utilizes an iterable dataset so that the training data is never fully loaded into memory at one time to prevent out-of-memory errors. The sequences are one-hot encoded with padding extending to the length of the longest sequence across the dataset. In alignment with the example script, the same hyper-parameters were used without performing hyper-parameter optimization. As in the MicrobeRT training pipeline, cross-entropy loss was used with class weighting and AdamW optimizer with weight decay [16].

#### Parameter-Efficient Fine-Tuning

Parameter-efficient fine-tuning (PEFT) methods have been increasingly used to reduce the memory requirements of fine-tuning large pre-trained language models by updating only a small subset of parameters while keeping the base model fixed. Common approaches, such as low-rank adaptation (LoRA) and infused adapter by inhibiting and amplifying inner activations ($(\text{IA})^3$), have been adapted to genomic language models (gLMs) [17, 18, 19, 20, 21].

In prior work [22], we systematically evaluated PEFT methods in a hierarchical classification setting with an expanding label space. While prior studies primarily focused on binary tasks (e.g., promoter/enhancer detection and signal peptide prediction), our results showed that PEFT performance degrades relative to full fine-tuning as the classification space grows, without clear gains in runtime or memory efficiency.

#### Performance Metrics

Classification tasks were evaluated using multiple complementary metrics. Due to the high-class imbalance across taxonomic classes, the macro-averaged F1 score (F1 Macro) was used as the primary measure for taxonomy models, since it treats all classes equally and emphasizes minority classes that might otherwise be overlooked [14]. For comparative purposes, the micro-averaged F1 (F1 Micro), weighted F1 (F1 Weighted), and Mean Reciprocal Rank (MRR) were also reported to provide additional perspectives on performance. Because AMR and pathogenicity classification involved fewer classes, the difference between macro- and weighted-averaging was less pronounced. For pathogenicity, balanced accuracy was also included as an evaluation metric, reflecting the importance of identifying pathogenic sequences. In this context, false negatives are more costly than false positives, and balanced accuracy ensures that recall is given equal consideration across both classes.

The F1 Macro computes the F1 score for each class and averages them equally, making it robust to class imbalance. In contrast, F1 Micro aggregates true positives, false positives, and false negatives across all classes before calculating the F1, thereby weighting each sample equally and reflecting overall performance. F1 Weighted provides an intermediate perspective by computing per-class F1 scores and averaging them proportionally to class size, ensuring that frequent classes exert more influence while still accounting for minority classes. Mean Reciprocal Rank (MRR) evaluates ranking performance by averaging the reciprocal of the rank of the first correct prediction across queries, emphasizing how highly relevant predictions appear. Finally, balanced accuracy calculates

Table 6: Summary of experiments optimizing hyperparameters for HyenaDNA-1M predictions for taxonomic classification

| Batch Size | Learning Rate | Weight Decay | Warmup Ratio | Balanced Accuracy (Genus) |
|---|---|---|---|---|
| 512 | 4E-04 | 0.01 | 0.05 | 0.160 |
| 256 | 2E-04 | 0.01 | 0.05 | 0.145 |
| 256 | 2E-04 | 0 | 0 | 0.101 |
| 256 | 1E-04 | 0.01 | 0.05 | 0.0750 |
| 512 | 2E-04 | 0.01 | 0.05 | 0.0663 |

the mean recall across classes, offering a fairer assessment in imbalanced settings and highlighting sensitivity to the minority (pathogenic) class where recall is critical.

#### Taxonomy

In fine-tuning the HyenaDNA class of models for taxonomic classification, a dataset of 155,725,641 sequences was annotated across five taxonomic ranks: superkingdom, phylum, family, genus, and species. The number of unique labels at each rank was 4, 79, 940, 4,224, and 19,402, respectively. A hold-out set of 1,572,987 sequences was reserved for validation during training. The sequence lengths ranged from 500 – 1750 bp. The per-device batch size was set to the maximum permitted by GPU memory, and training was conducted on 8 NVIDIA H100 GPUs for up to 5 epochs. The weight decay was 0.01, the cosine learning rate scheduler had a 0.05 warmup ratio, and bf16 precision was used. To prevent overfitting, early stopping with a patience of three epochs was incorporated. Because the dataset was too large to fit into CPU or GPU memory, a streaming-based batched tokenization and training was adopted. Exploratory experiments using a 0.05% subset of the full database, split into training and test sets, were performed prior to full-scale training. These preliminary experiments were used to select the optimal learning rate (given the fixed batch size) by evaluating model performance on the hold-out test set (Table 6).

An additional experiment was performed to further explore fine-tuning a full taxonomic classifier in two stages, inspired by "curriculum learning" strategies. In the first stage, the model was trained for five epochs with classification layers for superkingdom, phylum, family, and genus. In the second stage, an additional classification head for species was added, the model was initialized with weights from the previously trained genus-level classifier, and training was continued for another five epochs. The motivation for this staged "pre-training" phase was to leverage hierarchical taxonomic information to support more effective downstream learning and to improve performance on the species-level classification task, which is particularly challenging due to the large number of classes involved. Ultimately, the results from the "curriculum learning" train run were not significantly different from just training on all five taxonomic ranks at the same time, as shown in Table 7.

Training curves were assessed across classification tasks throughout the process of fine-tuning our models (Figure 3). The same fine-tuning strategy was applied to NT and DNABERT, with the exception that training was performed on a further processed version of the taxonomic database. Because transformer model memory scale quadratically with sequence length, training on the original dataset (up to 1,750 bp) would have required over a month, making it impractical. To mitigate this, sequences were chunked to <300 bp, which increased the dataset to 614,410,126 sequences but reduced the memory footprint per batch, allowing training to finish in under two weeks. A hold-out set of 606,971 sequences was reserved for validation. Given the substantial computational cost, training was restricted to a single full run each for NT-50M and DNABERT-2 (Table 8).

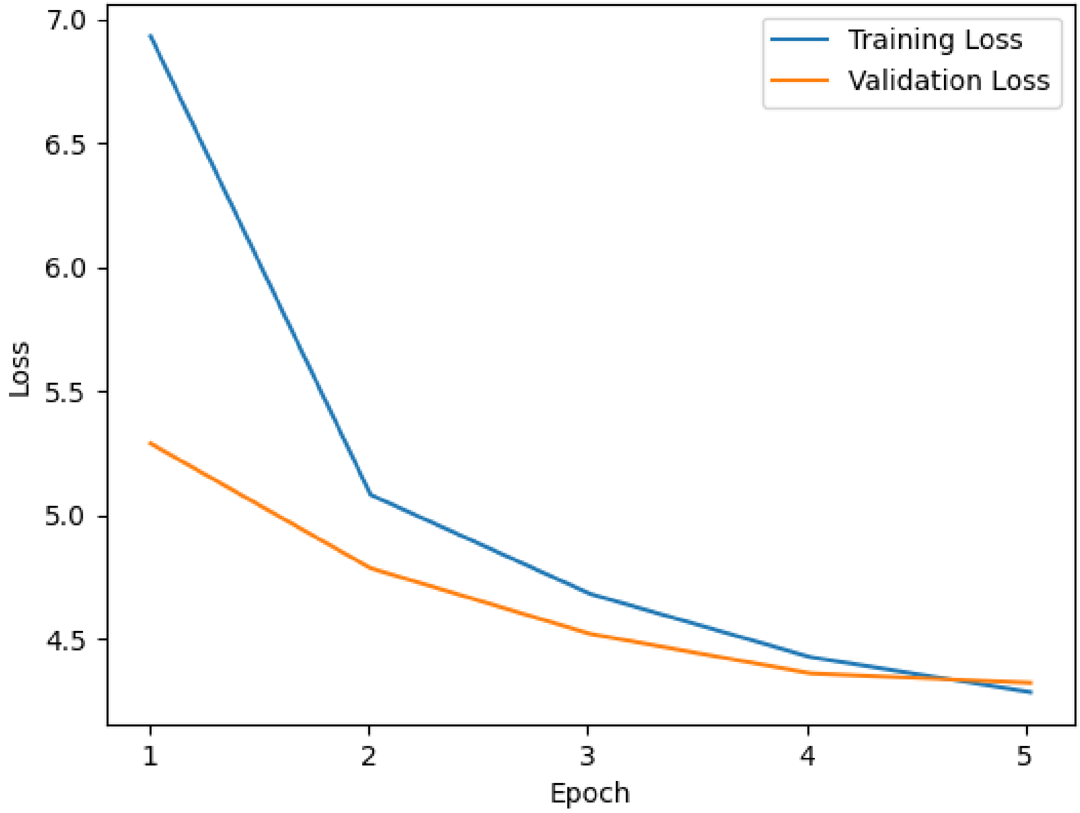


Figure 3: Example training history curve during fine-tuning of the DNABERT-2 model for taxonomic classification.

Table 7: Comparison of standard fine-tuning and curriculum learning for HyenaDNA models on taxonomic classification, showing limited performance gains from the two-stage curriculum learning approach.

| Model | Batch Size | Learning Rate | Training (hours) | Balanced Accuracy (Species) |
|---|---|---|---|---|
| hyenadna-large-1m-seqlen-hf | 512 | 0.0004 | 190.1 | 0.259 |
| hyenadna-medium-450k-seqlen-hf | 512 | 0.0004 | 180.9 | 0.263 |
| hyenadna-small-32k-seqlen-hf | 1024 | 0.0008 | 141.3 | 0.265 |
| hyenadna-large-1m-seqlen-hf (curriculum) | 512 | 0.0004 | 380 | 0.274 |

Table 8: Training hyperparameters and runtime for NT-50M and DNABERT-2 models fine-tuned on the taxonomic classification task.

| Model | Batch Size | Learning Rate | Training (hours) |
|---|---|---|---|
| nucleotide-transformer-v2-50m-multi-species | 2048 | 0.0005 | 260.9 |
| DNABERT-2-117M* | 4096 | 0.0001 | 299.4 |

* Trained over 4 GPUs rather than 8.

Table 9: Training hyperparameters and total wall-clock time for models trained on the binary antimicrobial resistance (AMR) classification dataset.

| Model | Batch Size | Learning Rate | Training (hours) |
|---|---|---|---|
| DNABERT-2-117M* | 920 | 2.68E-03 | 15.52 |
| DNABERT-S | 1840 | 3.87E-04 | 58.47 |
| hyenadna-large-1m-seqlen-hf | 6000 | 2.26E-03 | 36.13 |
| hyenadna-medium-160k-seqlen-hf | 6000 | 1.57E-03 | 36.39 |
| hyenadna-medium-450k-seqlen-hf | 6000 | 1.57E-03 | 32.63 |
| hyenadna-small-32k-seqlen-hf | 8000 | 3.90E-03 | 5.94 |
| nucleotide-transformer-v2-100m-multi-species* | 352 | 3.38E-04 | 86.74 |
| nucleotide-transformer-v2-250m-multi-species | 480 | 1.89E-04 | 76.03 |
| nucleotide-transformer-v2-50m-multi-species* | 600 | 7.53E-04 | 57.58 |
| Lyra** | 128 | 1e-3 | 57.12 |

* Trained over 4 GPUs rather than 8.

** Lyra used a single GPU at minimal capacity.

#### Antimicrobial Resistance

A substantial difference in training time between binary and multi-class classification tasks was observed during hyperparameter evaluation, as shown in Table 9 and Table 10. The consistently longer training times for Lyra compared to the other models are attributable to differences in computational resources: while the other models were trained using multiple GPUs, Lyra runs were limited to a single GPU with minimal utilization (at most 2 GB of VRAM).

Table 10: Training hyperparameters and total wall-clock time for models trained on the multi-class antimicrobial resistance (AMR) classification dataset.

| Model | Batch Size | Learning Rate | Training (hours) |
|---|---|---|---|
| DNABERT-2-117M | 1840 | 5.36E-03 | 0.135 |
| DNABERT-S | 1840 | 3.87E-04 | 0.135 |
| hyenadna-large-1m-seqlen-hf | 6000 | 2.26E-03 | 0.110 |
| hyenadna-medium-160k-seqlen-hf | 6000 | 1.57E-03 | 0.111 |
| hyenadna-medium-450k-seqlen-hf | 6000 | 1.57E-03 | 0.111 |
| hyenadna-small-32k-seqlen-hf | 8000 | 3.90E-03 | 0.090 |
| nucleotide-transformer-v2-100m-multi-species | 704 | 6.76E-04 | 0.111 |
| nucleotide-transformer-v2-250m-multi-species | 480 | 1.89E-04 | 0.148 |
| nucleotide-transformer-v2-50m-multi-species | 1200 | 1.51E-03 | 0.089 |
| Lyra** | 128 | 1e-3 | 0.25 |

** Lyra used a single GPU at minimal capacity.

#### Pathogenicity Classification

To classify pathogenicity, several of the previously discussed models were fine-tuned, including Lyra. As a comparator, an additional model specifically developed for the task of pathogenicity classification entitled Deep Learning Approach to Pathogenicity Classification (DeePaC [23]) was also fine-tuned.

The DeePaC model architecture is based on a reverse complement convolutional architecture. This, in principle, is similar to a bi-directional recurrent neural network, in which the sequence is passed through the model forward and through the same model again, but with the sequence reversed. This results in twice as many features as it passes through the model. DeePaC also does this but with convolutional neural networks, rather than recurrent neural networks, which are called reverse complement (RC) layers. The RC layers contain the functions for taking the input of a sequence, reversing it, passing the forward then the reversed sequences through the same convolution layer and concatenating them together, followed by a ReLU activation function. In the architecture used, which is the default from the paper, there is an initial dropout layer with a dropout rate of 0.25. The inputs are one-hot encodings of each base pair, so no embedding layer is used. The input is then fed into the first of two RC layers. Both RC layers, separated by a reverse complement pooling layer, both have filters of 512, kernel size of 15, and dilation and strides of 1. The reverse complement average pooling layer does an average pooling on the forward and reverse sequence separately. This is followed by a global average pooling layer and a RC dense layer. The RC dense layer has 256 units and uses a ReLU activation function that is applied to the forward and reverse components separately and concatenated. Then, a dense layer with 256 units is applied to the concatenated forward and reverse components before being applied to a dense layer with 1 unit for binary classification.

The models were all trained for 15 epochs with learning rates of $2e^{-}4$ using an Adam optimizer [16] and batch sizes of 512 (Table 11). As noted previously, Lyra runs only utilized a single GPU and at minimal capacity. As this is a binary classification problem, a binary cross entropy loss function is used.

In addition to the published fungal dataset ('DeePaC Fungal' [8]) and the simulated viral and bacterial datasets ('DeepSim Viral' and 'DeepSim Bacterial', simulated using DeepSimulator 1.5

Table 11: Training hyperparameters for models trained on the pathogenicity classification task.

| Model | Batch Size | Learning Rate |
|---|---|---|
| DNABERT-2-117M | 512 | 2e-4 |
| hyenadna-small-32k-seqlen-hf | 512 | 2e-4 |
| nucleotide-transformer-v2-50m-multi-species | 512 | 2e-4 |
| DeePaC | 512 | 2e-4 |
| Lyra* | 512 | 2e-4 |

* Lyra used a single GPU at minimal capacity.

Table 12: Training time required for each combination of model and dataset for pathogenicity classification

| Model | DeepSim Viral | DeepSim Bacterial | DeePaC Viral | DeePaC Fungal |
|---|---|---|---|---|
| DNABERT-2 | 2.67 h | 14.77 h | 22.8 h | 8 h |
| Hyenadna 32k | 1.62 h | 12.2 h | 8.25 h | 7.88 h |
| NT 50m | 1.18 h | 11.93 h | 8.03 h | 13 h |
| DeePaC* | 5.58 h | 22.93 h | 9.18 h | 24.05 h |
| Lyra* | - | 57.12 h | 11.63 h | 12.38 h |

* DeePaC and Lyra were trained using a single GPU; all others used 4 GPUs.

[9, 10]), an additional viral dataset ('DeePaC viral' [24]) was used for training and evaluation of the pathogenicity models (Table 12). Specifically, the nonpathogenic_train_strat (nonhuman pathogen data stratified in equal proportions chordate-infecting, metazoan-infecting, eukaryote-infecting, and all-nonhuman viruses) and pathogenic_train_hum files were used for training, and nonpathogenic_test_strat_1 and pathogenic_humspec_test_1 were used for testing. For validation, the nonpathogenic_val_strata and pathogenic_val_hum were used.

For DeepSim Bacterial and DeePaC fungal, the training datasets were sampled down to 20% of the original dataset size using random sampling due to memory constraints during training. Details of the training, testing, and validation sets are shown in Table 13.

Table 13: Train, test, and validation details for datasets prepared for pathogenic classification

| Dataset | Purpose | # Sequences | Fraction Pathogenic | Lengths (nt) |
|---|---|---|---|---|
| *DeePaC Viral* | | | | |
| | Training | 20,003,716 | 49.9% | 250 |
| | Testing | 1,250,465 | 49.9% | 250 |
| | Validation | 2,500,460 | 49.9% | 250 |
| *DeepSim Viral* | | | | |
| | Training | 2,260,207 | 66.5% | 140–950, mean 881 |
| | Testing | 282,716 | 66.5% | 140–950, mean 881 |
| | Validation | 282,717 | 66.5% | 140–950, mean 881 |
| *DeepSim Bacterial* | | | | |
| | Training | 12,475,872 | 52.0% | 140–950, mean 908 |
| | Testing | 6,238,628 | 51.9% | 140–950, mean 908 |
| | Validation | 6,238,628 | 51.9% | 140–950, mean 909 |
| *DeePaC Fungal* | | | | |
| | Training | 8,453,716 | 26.5% | 140–950, mean 867 |
| | Testing | 4,226,846 | 26.5% | 140–950, mean 867 |
| | Validation | 4,226,847 | 26.5% | 140–950, mean 867 |

### References


[1] Blin, Kai. ncbi-genome-download, 2023.

[2] RefSeq: NCBI Reference Sequence Database, Accessed 2024.

[3] NCBI GenBank, Accessed 2024.

[4] NCBI AMR Resources, Accessed 2024.

[5] Brian P Alcock, William Huynh, Romeo Chalil, Keaton W Smith, Amogelang R Raphenya, Mateusz A Wlodarski, Arman Edalatmand, Aaron Petkau, Sohaib A Syed, Kara K Tsang, Sheridan J C Baker, Mugdha Dave, Madeline C McCarthy, Karyn M Mukiri, Jalees A Nasir, Bahar Golbon, Hamna Imtiaz, Xingjian Jiang, Komal Kaur, Megan Kwong, Zi Cheng Liang, Keyu C Niu, Prabakar Shan, Jasmine Y J Yang, Kristen L Gray, Gemma R Hoad, Baofeng Jia, Timsy Bhando, Lindsey A Carfrae, Maya A Farha, Shawn French, Rodion Gordzevich, Kenneth Rachwalski, Megan M Tu, Emily Bordeleau, Damion Dooley, Emma Griffiths, Haley L Zubyk, Eric D Brown, Finlay Maguire, Robert G Beiko, William W L Hsiao, Fiona S L Brinkman, Gary Van Domselaar, and Andrew G McArthur. CARD 2023: expanded curation, support for machine learning, and resistome prediction at the Comprehensive Antibiotic Resistance Database. *Nucleic Acids Research*, 51(D1):D690–D699, January 2023.

[6] Martin Steinegger and Johannes Söding. Mmseqs2 enables sensitive protein sequence searching for the analysis of massive data sets. *Nature Biotechnology*, 35(11):1026–1028, 2017.

[7] Gustavo Arango-Argoty, Emily Garner, Amy Pruden, Lenwood S. Heath, Peter Vikesland, and Liqing Zhang. DeepARG: a deep learning approach for predicting antibiotic resistance genes from metagenomic data. *Microbiome*, 6(1):23, February 2018.

[8] Jakub M Bartoszewicz, Ferdous Nasri, Melania Nowicka, and Bernhard Y Renard. Detecting DNA of novel fungal pathogens using ResNets and a curated fungi-hosts data collection. *Bioinformatics*, 38(Supplement_2):ii168–ii174, 2022.

[14] Zachary Chase Lipton, Charles Elkan, and Balakrishnan Narayanaswamy. Thresholding Classifiers to Maximize F1 Score, 2014.

[15] Krithik Ramesh, Sameed M. Siddiqui, Albert Gu, Michael D. Mitzenmacher, and Pardis C. Sabeti. Lyra: An Efficient and Expressive Subquadratic Architecture for Modeling Biological Sequences, March 2025. arXiv:2503.16351 [cs].

[16] Ilya Loshchilov and Frank Hutter. Decoupled Weight Decay Regularization, 2017.

[17] Zeyu Han, Chao Gao, Jinyang Liu, Jeff Zhang, and Sai Qian Zhang. Parameter-Efficient Fine-Tuning for Large Models: A Comprehensive Survey, September 2024.

[18] Shuai Zeng, Duolin Wang, Lei Jiang, and Dong Xu. Parameter-efficient fine-tuning on large protein language models improves signal peptide prediction. *Genome Research*, 34(9):1445–1454, October 2024.

[19] Edward J. Hu, Yelong Shen, Phillip Wallis, Zeyuan Allen-Zhu, Yuanzhi Li, Shean Wang, Lu Wang, and Weizhu Chen. LoRA: Low-Rank Adaptation of Large Language Models, October 2021.

[20] Haokun Liu, Derek Tam, Mohammed Muqeeth, Jay Mohta, Tenghao Huang, Mohit Bansal, and Colin Raffel. Few-Shot Parameter-Efficient Fine-Tuning is Better and Cheaper than In-Context Learning, August 2022.

[21] Samuel Sledzieski, Meghana Kshirsagar, Minkyung Baek, Rahul Dodhia, Juan Lavista Ferres, and Bonnie Berger. Democratizing protein language models with parameter-efficient fine-tuning. *Proceedings of the National Academy of Sciences*, 121(26):e2405840121, June 2024.

[22] Daniel Berman, Daniel Jimenez, Stanley Ta, Brian Merritt, Jeremy D. Ratcliff, and Vijay Narayan. Evaluating the effectiveness of parameter-efficient fine-tuning in genomic classification tasks. *bioRxiv*, 2025. Preprint.

[23] Jakub M Bartoszewicz, Anja Seidel, Robert Rentzsch, and Bernhard Y Renard. Deepac: predicting pathogenic potential of novel dna with reverse-complement neural networks. *Bioinformatics*, 36(1):81–89, 2019.

[24] Bartoszewicz, Jakub M and Seidel, Anja and Renard, Bernhard Y. Interpretable detection of novel human viruses from genome sequencing data. *NAR Genomics and Bioinformatics*, 3(1), 2021.
